# Predicting near-future deforestation in West African Key Biodiversity Areas to inform conservation urgency

**DOI:** 10.1101/2024.10.07.616969

**Authors:** Brittany T. Trew, Graeme M. Buchanan, Felicity A. Edwards, Fiona J. Sanderson

## Abstract

Site-based protection is a cornerstone of 21^st^ century conservation and a core component of global biodiversity conservation targets. However, loss of tropical forests, the most biodiverse of habitats, is a major threat to such sites. Here, we predict near-future deforestation risk in 113 Key Biodiversity Areas (KBAs) - sites of objectively defined global conservation importance - in the Guinean Forest biodiversity hotspot and identify those factors associated with loss. We find that (i) KBAs in the Guinean Forests have lost over 265,000 hectares of forest cover over the past decade, with those in Côte d’Ivoire experiencing the highest forest loss; (ii) future deforestation risk is highest among KBAs in Liberia and Côte d’Ivoire (on average 10% predicted loss across KBAs), where some are predicted to lose over a fifth of remaining forest cover by 2033. Models indicate that deforestation is highly contagious, with historical forest loss effectively predicting further adjacent loss, and that forest fragmentation and ease of human access also increased the localised deforestation risk. Conversely, predicted forest loss was lower in sites under some form of conservation protection. Our methods to predict near-future hotspots of deforestation risk in KBAs are reproducible and therefore applicable to other biodiversity hotspots. In the Guinean forests, our results highlight where conservation interventions to mitigate forest loss should be urgently prioritised.

## 1.0 Introduction

Tropical forests support the majority of terrestrial biodiversity (Barlow et al., 2018; Pillay et al., 2022), providing critical ecosystem services and sustaining the livelihoods of over a billion people worldwide (Fedele et al., 2021). Despite their ecological significance, there have been substantial declines in tropical forest cover in recent decades (FAO & UNEP, 2020; Gibbs et al., 2010), as well as degradation of remaining habitat. The change in forest cover has resulted in widespread fragmentation (Ma et al., 2023) that not only compromises the integrity and resilience of these ecosystems (IPBES, 2019) but also precipitates further biodiversity loss (Alroy, 2017). This is of particular concern where it occurs in locations such as biodiversity hotspots (Myers et al., 2000) which host high concentrations of endemic species (Brooks et al., 2002, Barlow et al., 2018).

The Guinean Forests of West Africa is one such globally important biodiversity hotspot (Myers et al. 2000) and some of the most intact tropical forests in Africa (Crowe et al., 2023). The region supports 900 species of bird and almost 400 terrestrial mammal species, as well as providing vital ecosystem services to local communities and acting as an important source of identity and culture (CEPF, 2017). Its importance is reflected in the identification of over 100 Key Biodiversity Areas (KBAs) in the region (KBA Partnership, 2024). These sites are of global importance for their significant contribution to biodiversity persistence (IUCN, 2016) and an important tool for an evidence-based approach to expand site-based global conservation efforts in line with international ambition (Plumptre et al 2024). Nevertheless, KBAs are also acutely affected by deforestation (Tracewski et al., 2016), with a four-fold increase in deforestation rates detected across the hotspot between 2014 and 2019 (FAO, 2020) signalling an escalating crisis that threatens the very essence of their ecological significance.

Understanding factors that affect the risk of deforestation and forest degradation in this region is essential, both to prioritise conservation action, and for devising effective conservation strategies to safeguard these critical ecosystems. An extensive body of literature examines potential factors affecting the likelihood of pantropical forest loss and other conservation outcomes (e.g., Aide et al., 2013; Busch & Ferretti-Gallon, 2017; Geist & Lambin, 2002). In a recent review of global deforestation (Busch & Ferretti-Gallon, 2023), the number of deforestation events was found to be consistently higher (i) at lower elevations, (ii) on flatter slopes, (iii) on soil more suitable for agriculture, (iv) near ongoing clearing activity, and (v) near roads and urban areas. Deforestation in tropical countries has been found to be primarily driven by agricultural and urban expansion, infrastructure development and logging (Armenteras et al., 2017; Buchanan et al. 2009; Curtis et al., 2018; Gibbs et al., 2010), with the expansion of cocoa cultivation alone leading to the loss of millions of hectares of tropical forest in West Africa (Gockowski & Sonwa,□2011). Additionally, forest degradation - commonly defined as a reduction in forest biomass without changing land use (Thompson et al., 2013) - has been found to be non-randomly distributed (Matricardi□et al., 2020), corresponding to agricultural frontiers with high rates of deforestation, habitat fragmentation and infrastructure development (Bourgoin et al., 2021; Hansen et al., 2020; Tyukavina □et al., □2016).

The conservation and restoration of forests will not only preserve biodiversity but will also play a crucial role in mitigating the impacts of climate change in the coming decades (Goldstein et al., 2020, Trew et al., 2024a). Forests serve as significant carbon sinks, storing carbon in above- and below-ground biomass (Harris et al., 2021; Maxwell et al., 2019), and KBAs in west Africa store a volume of carbon greater than expected based on their area (Buchanan et al., 2021). Ergo, protecting and restoring high-integrity forest areas contributes to carbon sequestration, aiding global efforts to limit temperature rises and combat climate change. Recent pledges at international fora, such as the United Nations Framework Convention on Climate Change (UNFCCC), underscore the imperative to halt deforestation and restore tree cover by 2030, and specifically aim to halt forest loss in KBAs immediately.

This paper quantifies the spatiotemporal patterns of deforestation within KBAs in the Guinean Forests biodiversity hotspot. By employing a stepwise machine-learning modelling approach, we aim to predict hotspots of immediate deforestation risk to identify urgent conservation priorities and highlight potential drivers of deforestation that can be used to inform policy and management strategies in the region. We utilise a high resolution, historical forest cover dataset (Vancutsem et al., 2021) to analyse recent changes in forest cover across the KBAs and then identify those KBAs most at risk from deforestation using a dynamic and localised modelling methodology.

## 2.0 Methods

### 2.1 Historical forest cover change within the study region

The 621,705 km^2^ Guinean Forests hotspot spans nine West African countries and covers the Upper and Lower Guinean forests and four islands in the Gulf of Guinea, including lowland and montane forests and a wide range of habitats such as freshwater swamps, coastal habitats and wetland areas (CEPF, 2017). Annual data on the extent of tropical moist forest (TMF) cover for the hotspot was extracted from data prepared by the European Commission’s Joint Research Centre (Vancutsem et al., 2021) at the native 30 m gridded resolution and analysed in R version 4.2 (R Core Team, 2023). We considered all TMF cover changes that occurred between the start of 2013 and the end of 2023 - the most recent year for which data were available. Historical forest loss for the entire region was calculated as the difference between TMF forest cover in 2023 and TMF cover in 2013, inclusive of both undisturbed and degraded tropical moist forest. Forests in Togo and Benin were not included in any KBA analysis because the model groups here did not perform sufficiently.

### 2.2 Model training and testing

KBA boundaries (Birdlife International, 2023) were clipped to the boundary of the Guinean Forest hotspot (Hoffman et al, 2016). All KBAs located within the region were then grouped using k-means clustering by minimising the distance between individual KBA boundaries and the resulting centroid of each KBA group. This clustering method produced 30 groups of KBAs, henceforth model groups, which were filtered based on subsequent model testing (Fig. 1).

**Fig. 1.**
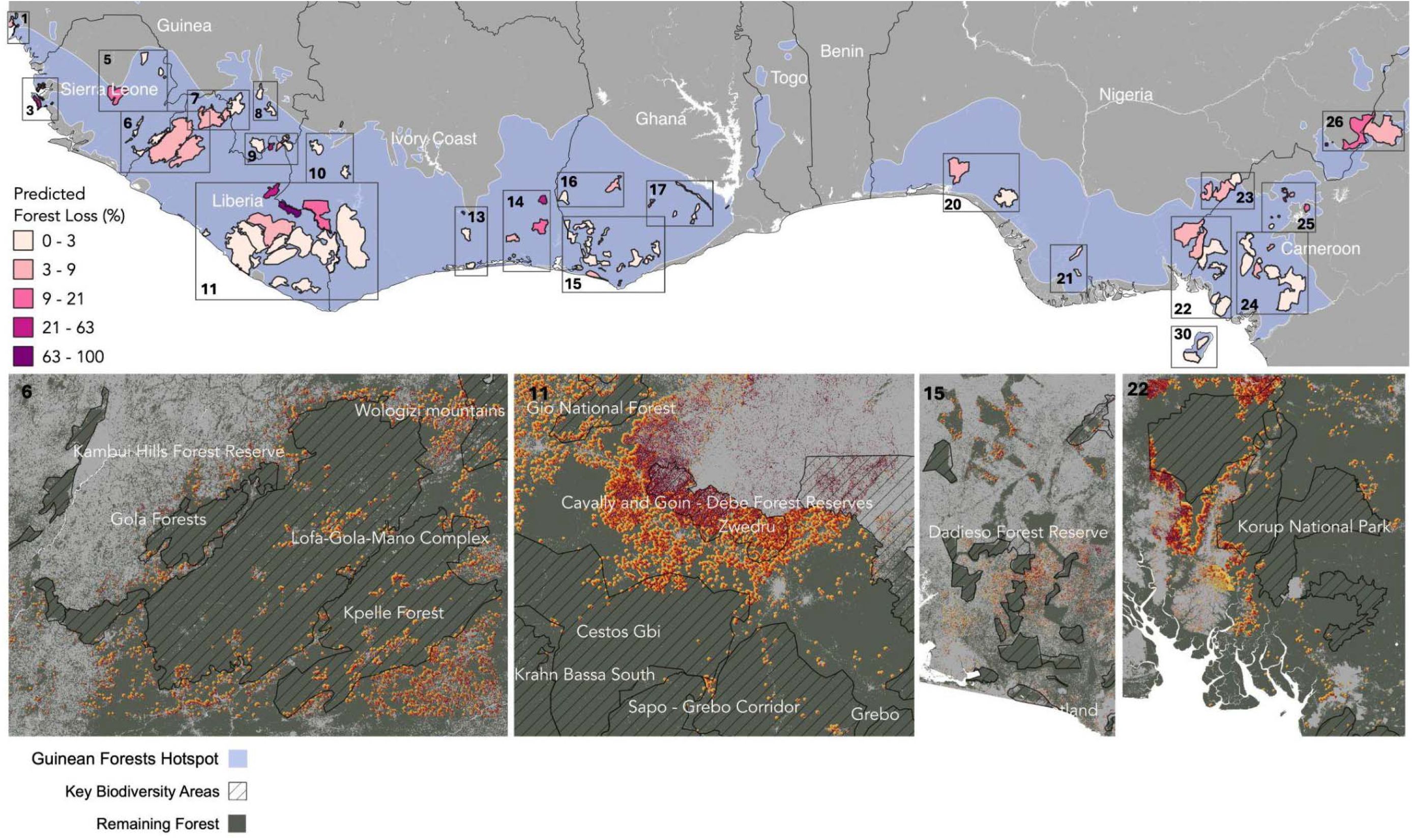
Map of the Guinean Forests of West Africa biodiversity hotspot (study extent), showing predicted forest loss (%) for each Key Biodiversity Area between 2025 and 2033. Inset maps show the projected year of deforestation up to 2033 for group 6 (Greater Gola Region on Sierra Leone/Liberia border), group 11 (Zwedru and surrounding KBAs in central Liberia), group 15 (KBAs in the southwestern lowland forests of Ghana), and group 22 (Cross River and Korup NP bordering Nigeria and Cameroon). See supplementary information (Fig. S3) for all group results.

For each of the KBA model groups, we built random forest regression models to predict the localised spatial pattern of deforestation risk. To build the model, we produced a training and testing detection/non-detection matrix of deforestation events for 2022 to 2023 (inclusive), using the most recent data on annual deforestation events from the TMF data (Vancutsem et al., 2021). The data were aggregated from 30 to 60 m gridded resolution using a thresholding method whereby a cell was deemed a deforestation event if at least 50% of the interior 30 m grid cells recorded a deforestation event. Model groups with less than 250 deforestation occurrence events for 2022 to 2023 were removed prior to modelling. Additionally, a set of explanatory variables was produced for 60 m gridded resolution, with the assumption that the environment two years prior to the detection/non-detection data (2020 to 2021) would impact the occurrence of deforestation events in the immediate aftermath (Boakes et al., 2009).

Explanatory variables were separated into static and dynamic variables (Supplementary Fig. S1). Static variables represented features assumed to stay constant through time, either due to data availability or the nature of the variable, while dynamic variables were re-calculated at each forward modelling step based on projected forest cover change. Explanatory variables were chosen to represent (i) landscape configuration - including forest fragmentation and connectivity metrics - (ii) forest access, (iii) anthropic influences such as the extent of neighbouring deforestation and degradation, local trends in forest clearance and conservation initiatives, and (iv) relevant natural features (see Table 1 for the full list of predictors considered for modelling). For each model group, to avoid strongly correlated predictors, we rejected variables with correlation problems by testing for multicollinearity, first using pairwise correlation (maximum threshold of 0.6) and then variance inflation analysis (maximum threshold of 5); whereby one of each variable pairing exceeding the correlation thresholds at each step was dropped. The combination of predictors chosen as best fit for each group are listed in supplementary Table S2.

**Table 1.** Full list of explanatory variables considered for the deforestation models (n = 13). For each individual KBA group, correlated explanatory variables were removed before model training and testing. For details regarding how these variables were derived, please see the supplementary methods.

| Category | Name | Description | Value to Model |
| --- | --- | --- | --- |
| Socioeconomic Influences | Deforestation anomaly | The recent 2-year anomaly in the mean number of deforestation events over the last 10 years within a local administrative area. | Captures any recent changes in deforestation rates between different administrative areas which can indicate changes in local attitudes towards forest loss as trends in the number of deforestation events change. |
| Forest Access | Road density | The length of all roads (m) within a moving window around a forest cell. | The proximity to roads and urban areas has consistently been found to be associated with a higher number of deforestation (Busch & Ferretti-Gallon, 2023). Additionally, degradation has been found to be higher near infrastructure development (Tyukavina et al., 2016). |
| Forest Access | Cost of Motor Access | Relative distance cost to the nearest village, town or city by motor vehicle from a forest cell. | Indicates the ease of access to forest using a motor vehicle. Accessibility has consistently been found to be associated with deforestation (Busch & Ferretti-Gallon, 2023) |
| Natural Features | Elevation | Elevation bands, every 100 m. | Higher elevations have been shown to consistently be associated with lower deforestation rates (Busch & Ferretti-Gallon, 2023). |
| Natural Features | Slope | Mean slope to the horizon (0-360 degrees). | Steeper slopes have been shown to consistently be associated with lower numbers of deforestation events (Busch & Ferretti-Gallon, 2023). |
| Forest Landscape | Euclidean Nearest Neighbour Distance | The ENN distance measures the distance of a forest cell to the next nearest forest cell and thus it describes forest patch isolation within the wider landscape. | Small forest fragments are more likely to disappear completely because the creation of new forest edges (edge effects) increases accessibility (Angelsen & Kaimowitz, 1999; Laurence et al., 2002; Pfaff, 1999), as well as tree mortality and fire frequency (Alves 2002; Broadbend et al., 2008). Decreasing fragment size has been specifically linked to tropical forest degradation (Bourgoin et al., 2021; Hansen et al., 2020). |
| Forest Landscape | Core Area Index | The CAI measures the percentage of a forest patch which is considered a core area. Core areas only include those forest grid cells which are exclusively surrounded by other forest cells. | As above. |
| Forest Landscape | Radius of Gyration | The radius of gyration measures the dispersion of forest patches within the landscape. This metric indicates the level of fragmentation at the landscape scale. | As above. |
| Forest Landscape | Perimeter-area ratio | The perimeter-area ratio is a measure of patch complexity, whereby the higher the perimeter length compared to the internal area, the more complex the forest patch shape. | As above. |
| Forest Landscape | Contiguity Index | The contiguity index assesses the spatial connectedness of forest grid cells within each forest patch. | As above. |
| Socioeconomic Influences | Neighbouring deforestation | Proportion of surrounding forest which was recently deforested (%) | The proportion of surrounding forest which was recently deforested has been used in other studies to model deforestation (Silva et al., 2022, Rosa et al., 2013). Deforestation has been found to occur primarily near previously deforested areas (Alves, 2002; Aguiar et al., 2007) and the number of deforestation events was found to be consistently lower away from ongoing clearing activity (Busch & Ferretti-Gallon, 2023). |
| Socioeconomic Influences | Neighbouring degradation | Proportion of surrounding forest which was recently degraded (%) | Degradation has been found to correspond to agricultural frontiers with high rates of deforestation, habitat fragmentation and infrastructure development (Tyukavina et al., 2016). Degraded forests with more than 50% canopy loss are significantly more vulnerable to subsequent deforestation (Bourgoin et al., 2024). |
| Socioeconomic Influences | Protected Area Designation | Protected area designation using IUCN categories (I-VI). | Powlen et al., 2023 found that PA placement had substantial influence over forest loss. Lower levels of deforestation were also found to be associated with forests under community management and African forests in protected areas (Bowker et al., 2017). |

Random forest regression has been found to be more accurate and consistent than random forest classification (Malley et al., 2012) and can be used to examine a large number of predictor variables simultaneously, while adjusting for multicollinearity, higher order interactions and nonlinear relationships (Breiman, 2001). As a result, it has been used successfully to investigate nonlinear drivers of forest loss (Aide et al., 2012; Zanella et al., 2017). However, spatial autocorrelation effects need to be mitigated and so, before training, each model group was divided into ten spatially disaggregated blocks (*sensu* Yates et al., 2023) and grid cells were all assigned a block number (1-10) for subsampling. Moreover, though random forest models are inherently less prone to overfitting than other methods, in highly complex mosaic landscapes such as this, overfitting to the training data may occur to some degree. To minimise this, we fit a down-sampled random forest model to the data (Wright, 2017; R Core Team, 2023), to balance the validation data, using an equal number of detections and non-detections (deforestation events versus tropical forest). Additionally, tree depth and node number were constrained.

A model was built for each combination of the spatially disaggregated blocks, whereby nine blocks were used to build the model and the tenth omitted block was used for testing. For each model block, the estimated occurrence rate of deforestation events was compared to the actual occurrence rate using a logarithmic loss performance metric (Yates et al., 2023). The relationship between the resulting predicted probability of a deforestation event and observed detection/non-detection of deforestation events was then fitted using a generalised additive model that was shape constrained to be monotonically increasing, using the *scam* package (Pya & Wood, 2015; R Core Team, 2023) and assessed using area under the curve (AUC); see supplementary Results and Table S3 for model evaluation scores.

The same model building process was repeated to train and test a degradation (defined by Vancutsem et al., 2021) model for each group, using annual remote-sensed observations of TMF degradation events (Vancutsem et al., 2021). Predictor variables were consistent with the deforestation models, except the recent degradation anomaly was used in place of recent deforestation anomaly.

To understand how explanatory variables affected the models, the permutation importance of each variable was calculated for each model group across all ten models (supplementary Table S4). We chose permutation over impurity-based importance as the latter is known to inflate the importance of numerical features (Breiman, 2001). A final weighted mean importance value for each predictor was calculated based on the same weightings used for the ensemble model. Additionally, to better understand the direction of the relationships between the variables of highest importance and forest loss, we extracted the combined effects values to derive accumulated local effects (ALE) plots for each variable across all models within all groups.

### 2.3 Predicting future deforestation risk

To project deforestation up to 2033, we used a stepwise forward modelling process (supplementary Fig. S1). For each model group, the probability of occurrence for both deforestation and degradation events was modelled for 2024 to 2025 (inclusive). Probabilities were projected individually using all the trained models (n=10) and explanatory variables for 2022 to 2023 for each model group. To account for confidence levels across the trained models, we derived a weighted ensemble mean probability of deforestation - as well as for degradation - for each model group, using the area under the curve (AUC) scores of each calibrated model as a weighting. Additionally, we calculated the coefficient of variation across trained models for each group, where low values imply high agreement between models.

To derive a threshold for classifying the continuous output from the models into a binary output of whether a deforestation or degradation event is projected to occur, a historical sensitivity analysis was performed for each model group (see supplementary Methods for further detail). These binary projections of events or non-events were then used to derive new forest cover maps for the start of 2026 and ergo new dynamic explanatory variables as per the methods. Projections of deforestation and degradation events for 2026 to 2027 (inclusive) were derived as above and this process was repeated every two years up to the end of 2033.

## 3.0 Results

### 3.1 Historical forest cover change

Approximately 17%, over 4.5 million hectares, of tropical forest in the Guinean Forests hotspot was lost between the start of 2013 and the end of 2023. Côte d’Ivoire lost the greatest proportion of remaining forest (∼ 34% and over 1.5 million hectares) to deforestation in the same period (Supplementary table S5). Sierra Leone, Ghana and Guinea were also badly affected, all losing over 20% of remaining forest cover and over 1.6 million hectares between them. Deforestation within KBAs in the Guinean Forests was generally lower than outside of their boundaries, but 265,00 hectares of forest was lost inside of KBAs in the same period. Some KBAs, particularly those located in Côte d’Ivoire and Sierra Leone, were badly affected by forest loss, losing over 10% of forest cover on average (Supplementary table S6). For instance, the Cavally and Goin-Debe FRs lost over 80,000 hectares of forest (over 52% of forest extent). Nevertheless, a quarter of KBAs in the region lost less than 1% of forest in the same period, including Mount Nlonako in Cameroon (0.28% loss), and the Cross River NP in Nigeria (0.92% loss).

### 3.2 Future deforestation hotspots

Almost a quarter of KBAs across the Guinean Forests hotspot were predicted to lose more than 5% of their current forest area (Fig. 1). The highest loss of forest by 2033 was projected for KBAs in Liberia and Côte d’Ivoire, where KBAs were predicted to lose approximately 10% of forest on average (Supplementary table S7). Specifically, Zwedru, Kpelle Forest, the Lofa-Gola-Mano Complex and the Wologizi Mountains were predicted to lose almost 100,000 hectares of forest between them (Supplementary table S8). Zwedru was particularly affected (Fig.2) and was predicted to lose over 48,300 hectares of forest (92% of the KBA’s current forest cover), as was the Cross River NP in Nigeria which was predicted to lose 24,548 hectares of forest. Additionally, several KBAs were all projected to lose over a fifth of their remaining forest cover, including Bossematie FR (63.2%) and the Lamto Ecological Research Station (38.33%) in Côte d’Ivoire, the Mbi Crater FR in Cameroon (47.5%), West Nimba in Liberia (34.8%), Western Area Peninsula and Yawri Bay in Sierra Leone (32.9% and 20.5% respectively) and Gio National Forest in Liberia (25.8%).

**Fig. 2.**
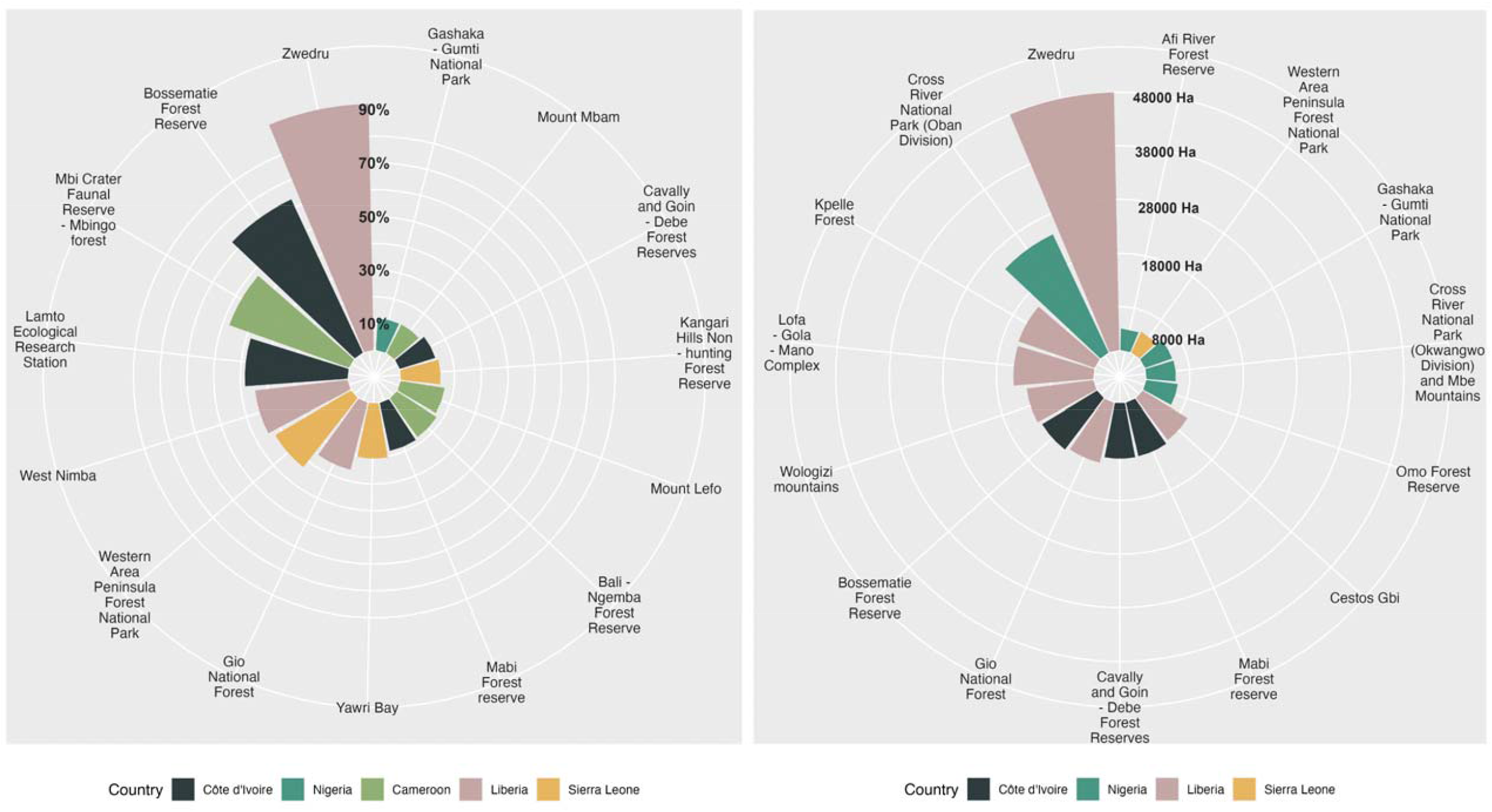
The total predicted loss (%) of currently remaining forest (observed in 2023) between 2025 and 2033 (inclusive) for Key Biodiversity Areas where predicted forest loss exceeded 10% and (B) KBAs with the highest total projected forest loss (Ha) between 2025 and 2033 (inclusive).

### 3.3 Correlates of deforestation risk

Neighbouring deforestation was the most important influence across model groups (Fig.3; supplementary Table S3). Forest accessibility and the recent anomaly in local deforestation rates were also particularly important across groups. Landscape configuration variables were often spatially correlated and therefore only used to train some model groups. Nevertheless, when used, the radius of gyration, measuring dispersion of forest patches across the landscape, substantially contributed to improving model performance.

**Fig. 3.**
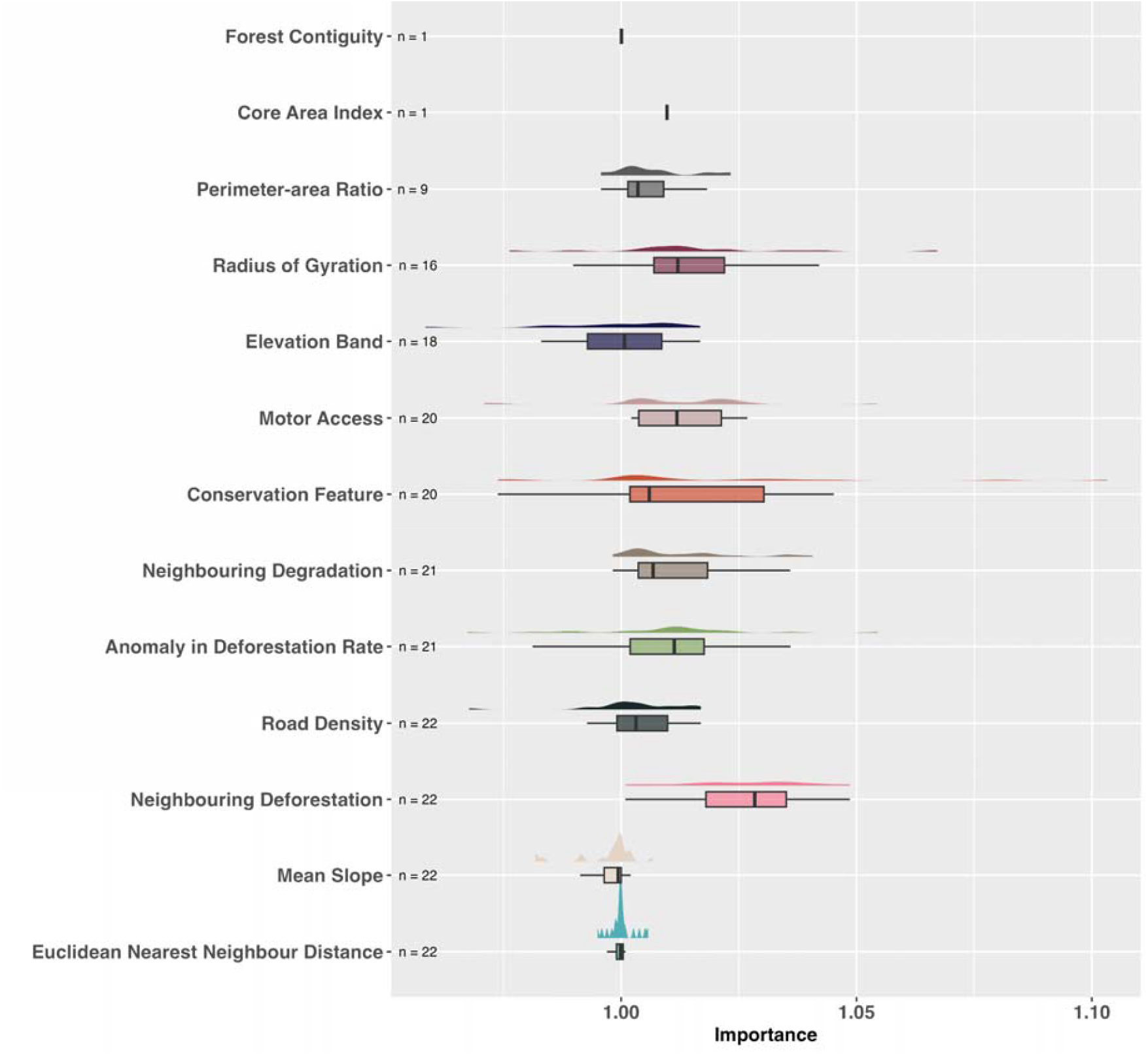
Distribution of the mean importance (increase in model performance) for each explanatory variable across all model groups (n=24). The observe distribution of importance values is plotted above each bar. Due to multicollinearity between some variables, model groups did not always use the entire set of predictor variables and so the number of times the variable was used is indicated by ‘n’. Please see supplementary Table S1 for a full table of variables used for model building by group. Importance values are an estimate of the level of influence and do not indicate direction of influence.

None of the relationships driving the models - those between the explanatory variables and deforestation events - were linear (Fig. 4; see supplementary Fig. S3 for individual group results). As expected, a greater proportion of neighbouring deforestation and degradation events increased the future risk of deforestation. Sharp increases in deforestation risk occurred in conjunction with low levels of neighbouring deforestation and degradation, as well as high road densities and increasing local deforestation rates (positive recent anomalies in the mean number of deforestation events within local administrations). Deforestation risk also typically increased when values of landscape configuration variables implied low forest connectivity and high fragmentation. For instance, highly fragmented forest, represented as a lower Euclidean nearest neighbour distance or radius of gyration, increased deforestation risk. In contrast, a higher cost to access forest for motor vehicles decreased the risk of deforestation, as did any protected area designation.

**Fig. 4.**
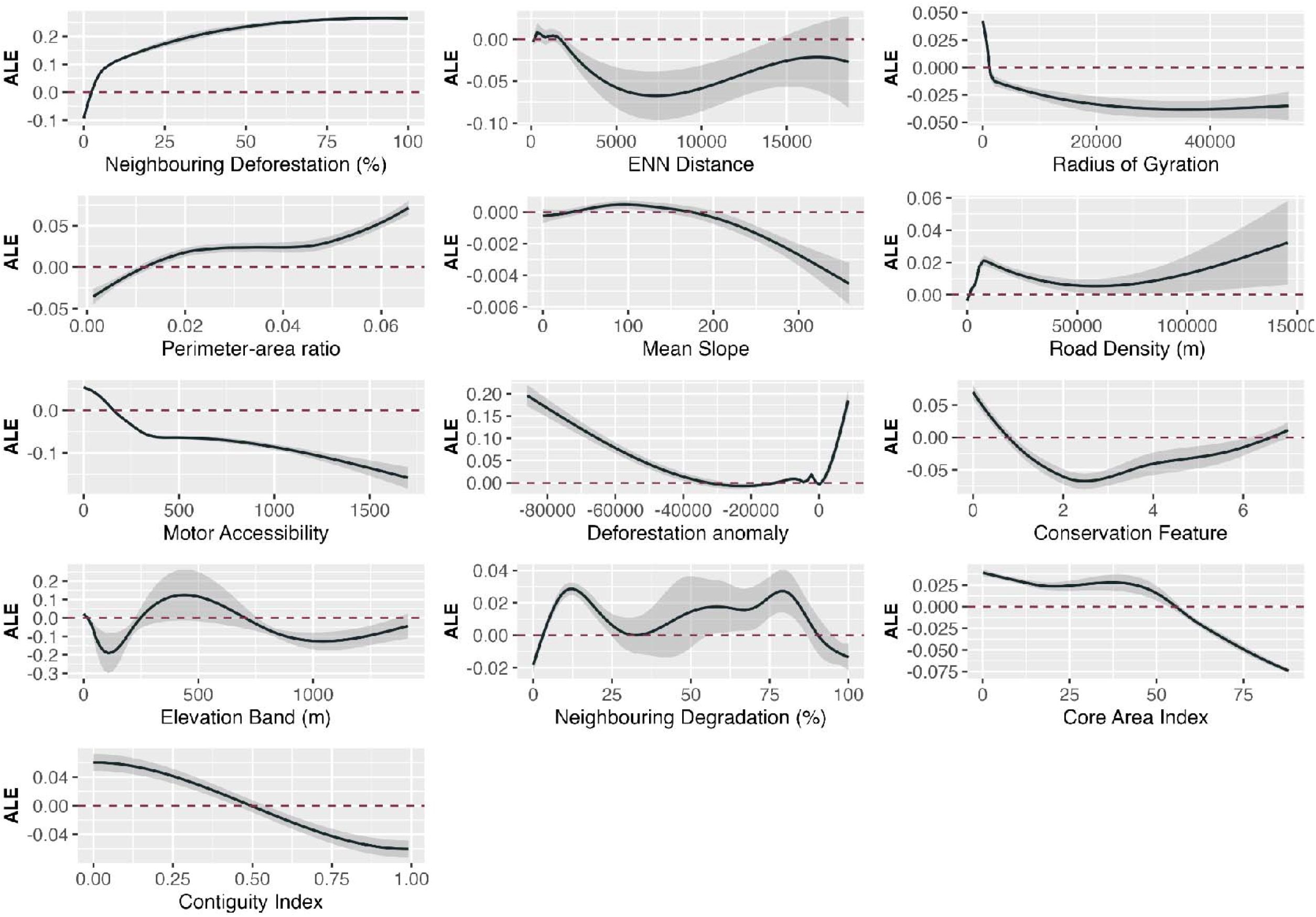
Accumulated local effects of predicted probability of deforestation and all explanatory variables used by the model groups. 95% confidence intervals displayed as greyed area. Conservation features are classed as: IUCN categories Ia & Ib = 1, II = 2, III = 3, IV = 4, V = 5, VI = 6 and None = 7.

Protected area designation generally lowered the risk of deforestation inside of PAs (Fig. 5); defined here as protected areas with IUCN categories I-IV. The difference between the probability of deforestation inside and outside of PAs was most pronounced in KBAs in Guinea, Côte d’Ivoire and Equatorial Guinea. However, formal protections did not lower risk in some regions. For instance, we found that areas of KBAs overlapping PAs in Nigeria were just as much at higher risk of near-future deforestation than those outside of PAs.

**Fig. 5.**
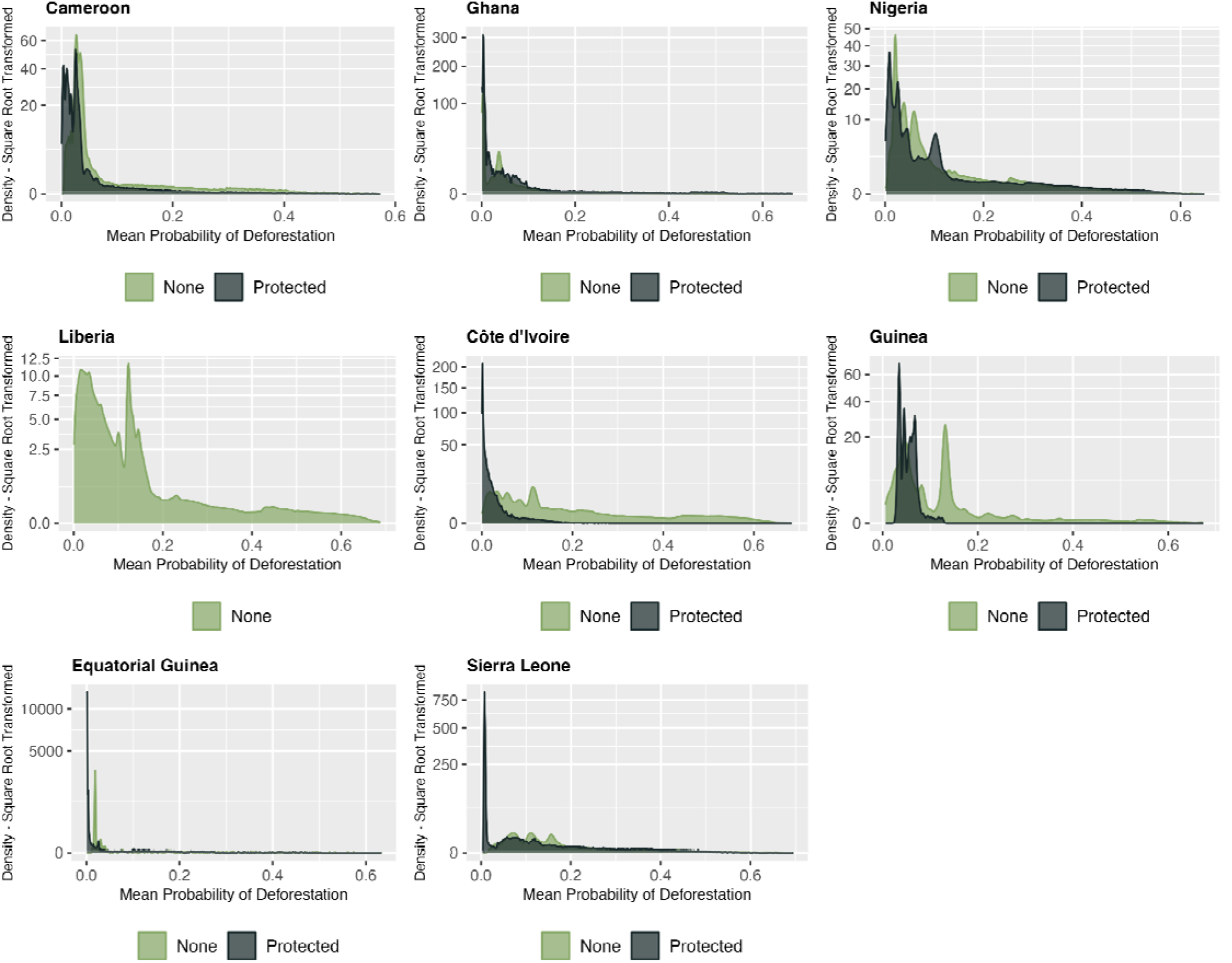
Distribution of the mean probability of deforestation between 2025 and 2033 (inclusive) for Key Biodiversity Areas which overlap protected areas (IUCN categories I-IV only), separated by forest location inside versus outside of these formal protections.

## 4.0 Discussion

We present the first near-future deforestation risk projections for KBAs in West Africa to better inform emerging global and regional conservation strategies. By training a stepwise random forest model on historical deforestation and degradation occurrence data, we have been able to predict dynamic high resolution spatial patterns of deforestation risk for all KBAs in the Guinean Forests of West Africa. Our findings reveal that KBAs in the hotspot have lost a substantial amount of forest cover between 2013 and 2023, with KBAs in Côte d’Ivoire losing the highest proportion of their remaining forest. Model predictions indicate that KBAs continue to face widespread and severe near-future risk from forest loss and fragmentation, especially in Liberia and Côte d’Ivoire, where some KBAs are projected to lose over a fifth of remaining forest cover by 2033. The continued rapid loss of forest from these areas threatens the entire hotspot region, as well as the ecological resilience, integrity, and ergo the biodiversity value of these KBAs.

Here, we assume temporally consistent relationships between explanatory variables and deforestation risk. However, factors associated with deforestation are naturally dynamic and thus there are inherent challenges and limitations to predicting where deforestation will occur next. As a result, the models are unable to account for changes to these relationships such as regional policy change, emerging market dynamics or unexpected climate events which can rapidly alter deforestation dynamics in ways that are difficult to predict. Moreover, the models are trained and tested on historical remotely sensed data which has inherent uncertainty. Nevertheless, as well as clearly demonstrating the complexity of relationships between factors associated with forest loss and its occurrence, where other studies have frequently oversimplified these relationships by using linear modelling approaches (Powlen et al., 2023), our methods allow us to derive a more accurate and dynamic risk analysis which can be updated annually, supporting urgent and targeted conservation decision-making.

Over the last decade, KBAs in the Guinean Forests hotspot have experienced less forest loss than those areas outside KBAs. Community-based forest management and programs focused on sustainable livelihoods, such as Reducing Emissions from Deforestation and Degradation (REDD+), have already led to positive forest conservation outcomes in the region and elsewhere (Agrawal et al., 2011; Guizar-Coutiño 2022; Holloway & Giandomenico, 2009; Jayachandran et al., 2017; Malan et al., 2024; Simonet et al., 2019). Many KBAs in the region also benefit from some form of protected area (PA) coverage, which generally lowered deforestation risk. However, our results demonstrate that gazetting does not remove risk entirely: forest loss persists in PAs, albeit often less than in unprotected areas (Geldmann et al., 2019; Wolf et al., 2021). We found considerable variation between countries, with deforestation risk generally higher in protected areas in Nigeria, Ghana and Sierra Leone. This may result from various factors, including how well conservation laws are enforced at country level (Geldmann et al., 2019).

The existence of roads within protected areas has been found to result in almost as much forest loss as outside them (Engert et al., 2024; Laurance et al., 2017). Limiting unregulated and undocumented road expansion within and around KBAs is an urgent priority, irrespective of PA coverage. Lower forest accessibility was associated with a steep decrease in projected deforestation risk and so, KBAs which are more remote are likely to be less at risk from deforestation (Simkins et al., 2024). Significant gaps in road mapping are not unusual, especially for developing nations (Kleinschroth et al., 2019; Engert et al., 2024) and we caution that our estimates of forest accessibility will be conservative and therefore deforestation risk may in some cases be underestimated. Moreover, steep mountain regions with high terrain complexity are inherently less accessible and will be at lower risk of deforestation.

KBAs may be intrinsically less vulnerable to forest loss in the Guinean Forests hotspot due to higher forest integrity (Crowe et al., 2023; Grantham et al., 2020). Here, we have used dynamic forest landscape metrics and contagion measures to represent elements of forest integrity (Grantham et al., 2020), such as edge effects, surrounding deforestation and degradation pressure and forest extent. These dynamic predictors were important for the deforestation-risk models, and measures indicating better forest integrity such as a higher forest perimeter-area ratio or lower neighbouring deforestation consistently preceded projections of lower deforestation risk to the KBA. Forest fragmentation is known to result in cascading deforestation effects (Busch & Ferretti-Gallon, 2023; Hansen et al., 2020) and so the continued forest integrity of KBAs in the Guinean Forests region is critical in minimising future forest loss.

We have identified those KBAs in the West African Guinean Forests hotspot which are at high risk of imminent widespread deforestation, where the implementation of evidence-based intervention to reduce future forest loss is urgent. Minimising future deforestation in KBAs is paramount in maintaining their high value and irreplaceability for biodiversity, carbon storage (Buchanan et al., 2020), the broader landscape’s ability to adapt to climate change through climate regulation and landscape connectivity (Trew et al., 2024a; Trew et al., 2024b). The conservation of KBAs in this region demands an integrated, multifaceted approach that recognizes the interdependencies between biodiversity conservation, socio-economic development, and climate mitigation (Gurney et al., 2023). Local stakeholder-informed area-based conservation measures, and social equity and community-based forest management programs, as well as better PA enforcement in some areas, are urgently needed for high-risk KBAs. Forest restoration, increasing forest integrity, in low-risk KBAs would add to their resilience to future deforestation risks. Our results underscore the urgent need for targeted conservation efforts across the West African region and the need for effective policy and management strategies to safeguard tropical biodiversity.

## Data Availability Statement

This study uses freely available data, as detailed in the supplementary methods, including: (1) the global tropical forest monitoring dataset (available at https://forobs.jrc.ec.europa.eu/TMF), (2) locations of roads and settlements obtained from Open Source Mapper (available at: https://www.openstreetmap.org), (3) river locations from World Wildlife Fund’s hydroshed database (available at: https://www.hydrosheds.org/products/hydrorivers), (4) the World Database on Protected Areas (available at: https://www.protectedplanet.net), (5) Key Biodiversity Area boundaries (available on request at: https://www.keybiodiversityareas.org/kba-data/request), (6) European Space Agency’s land cover data (available at: https://climate.esa.int/en/projects/land-cover/), (7) Digital elevation model from Amazon Web Services (available at: https://aws.amazon.com/public-datasets/terrain/), (8) Subnational administrative boundaries from the COD - Subnational Administrative Boundaries database (available at https://data.humdata.org/).

## Code Availability Statement

Code used for the analysis is available via Github at https://github.com/brittany-trew/Deforestation_WestAfrica with fully reproducible scripts to derive results shown here.

## Notes

**Conflict of Interest Statement** The authors declare no conflicts of interest.

### Competing Interest Statement

The authors have declared no competing interest.

## References

Agrawal, A., Nepstad, D., Chhatre, A. (2011) Reducing emissions from deforestation and forest degradation. Annual Review of Environment and Resources 36, 373–396.

Aide, T. M., Clark, M. L., Grau, H. R., López-Carr, D., Levy, M. A., Redo, D., … Muñiz, M. (2013). Deforestation and Reforestation of Latin America and the Caribbean (2001–2010). Biotropica, 45(2), 262–271.

Aguiar, A. P. D., Câmara, G., & Escada, M. I. S. (2007). Spatial statistical analysis of land-use determinants in the Brazilian Amazonia: Exploring intra-regional heterogeneity. Ecological Modelling, 209(2-4), 169–188.

Alroy, J. (2017) Effects of habitat disturbance on tropical forest biodiversity. Proceedings of the National Academy of Sciences 114, 6056–6061.

Alves, D. S. (2002). Space-time dynamics of deforestation in Brazilian Amazonia. International Journal of Remote Sensing, 23(14), 2903–2908.

Angelsen, A., & Kaimowitz, D. (1999). Rethinking the causes of deforestation: lessons from economic models. The world bank research observer, 14(1), 73–98.

Armenteras, D., Espelta, J. M., Rodríguez, N., & Retana, J. (2017). Deforestation dynamics and drivers in different forest types in Latin America: Three decades of studies (1980–2010). Global Environmental Change, 46, 139–147.

Barlow, J., França, F., Gardner, T. A., Hicks, C. C., Lennox, G. D., Berenguer, E., … Graham, N. A. J. (2018). The future of hyperdiverse tropical ecosystems. Nature, 559(7715), 517–526.

BirdLife International (2023). World Database of Key Biodiversity Areas (KBA): KBA Digital Boundaries. Developed by the KBA Partnership. Available at: https://www.keybiodiversityareas.org.

Breiman, L. (2001). Random Forests. Machine Learning, 45(1), 5–32.

Brooks, T.M., Mittermeier, R.A., Mittermeier, C.G., Da Fonseca, G.A.B., Rylands, A.B., Konstant, W.R., Flick, P., Pilgrim, J., Oldfield, S., Magin, G., Hilton-Taylor, C. (2002) Habitat Loss and Extinction in the Hotspots of Biodiversity. Conservation Biology 16, 909–923.

Boakes, E. H., Mace, G. M., McGowan, P. J. K., & Fuller, R. A. (2009). Extreme contagion in global habitat clearance. Proceedings of the Royal Society B: Biological Sciences, 277(1684), 1081–1085.

Bourgoin, C., Ceccherini, G., Girardello, M., Vancutsem, C., Avitabile, V., Beck, P. S. A., … Achard, F. (2024). Human degradation of tropical moist forests is greater than previously estimated. Nature.

Buchanan, G. M., Butchart, S. H. M., Chandler, G., & Gregory, R. D. (2020). Assessment of national-level progress towards elements of the Aichi Biodiversity Targets. Ecological Indicators, 116, 106497.

Buchanan, G. M., Field, R. H., Bradbury, R. B., Luraschi, B., & Vickery, J. A. (2021). The impact of tree loss on carbon management in West Africa. Carbon Management, 12(6), 623–633.

Busch, J., & Ferretti-Gallon, K. (2017). What Drives Deforestation and What Stops It? A Meta-Analysis. Review of Environmental Economics and Policy, 11(1).

Busch, J., & Ferretti-Gallon, K. (2023). What Drives and Stops Deforestation, Reforestation, and Forest Degradation? An Updated Meta-analysis. Review of Environmental Economics and Policy, 000–000.

Critical Ecosystem Partnership Fund (CEPF). 2017. Guinean Forests of West Africa Biodiversity Hotspot Ecosystem Profile. Washington, D.C.: CEPF. Available online: https://www.cepf.net/our-work/biodiversity-hotspots/guinean-forests-west-africa

Crowe, O., Beresford, A.E., Buchanan, G.M., Grantham, H.S., Simkins, A.T., Watson, J.E.M., Butchart, S.H.M. (2023) A global assessment of forest integrity within Key Biodiversity Areas. Biological Conservation 286, 110293.

Curtis, P. G., Slay, C. M., Harris, N. L., Tyukavina, A., & Hansen, M. C. (2018). Classifying drivers of global forest loss. Science, 361(6407), 1108–1111.

Engert, J.E., Campbell, M.J., Cinner, J.E., Ishida, Y., Sloan, S., Supriatna, J., Alamgir, M., Cislowski, J., Laurance, W.F. (2024) Ghost roads and the destruction of Asia-Pacific tropical forests. Nature.

Fedele, G., Donatti, C. I., Bornacelly, I., & Hole, D. G. (2021). Nature-dependent people: Mapping human direct use of nature for basic needs across the tropics. Global Environmental Change, 71, 102368.

Food and Agriculture Organization (FAO) & United Nations Environment Programme (UNEP). (2020). The state of the world’s forests: Forests, biodiversity, and people. Food and Agriculture Organization.

Farr, T.G., Rosen, P.A., Caro, E., Crippen, R., Duren, R., Hensley, S., Kobrick, M., Paller, M., Rodriguez, E., Roth, L., Seal, D., Shaffer, S., Shimada, J., Umland, J., Werner, M., Oskin, M., Burbank, D., and Alsdorf, D.E., 2007, The shuttle radar topography mission: Reviews of Geophysics, v. 45, no. 2, RG2004.

Geist, H. J., & Lambin, E. F. (2002). Proximate Causes and Underlying Driving Forces of Tropical Deforestation: Tropical forests are disappearing as the result of many pressures, both local and regional, acting in various combinations in different geographical locations. BioScience, 52(2), 143–150.

Geldmann, J., Manica, A., Burgess, N. D., Coad, L., & Balmford, A. (2019). A global-level assessment of the effectiveness of protected areas at resisting anthropogenic pressures. Proceedings of the National Academy of Sciences, 116(46), 23209–23215.

Gibbs, H. K., Ruesch, A. S., Achard, F., Clayton, M. K., Holmgren, P., Ramankutty, N., & Foley, J. A. (2010). Tropical forests were the primary sources of new agricultural land in the 1980s and 1990s. Proceedings of the National Academy of Sciences, 107(38), 16732–16737.

Gockowski, J., Sonwa, D. (2011) Cocoa Intensification Scenarios and Their Predicted Impact on CO2 Emissions, Biodiversity Conservation, and Rural Livelihoods in the Guinea Rain Forest of West Africa. Environmental Management 48, 307–321.

Goldstein, A., Turner, W. R., Spawn, S. A., Anderson-Teixeira, K. J., Cook-Patton, S., Fargione, J., … Hole, D. G. (2020). Protecting irrecoverable carbon in Earth’s ecosystems. Nature Climate Change, 10(4), 287–295.

Grantham, H.S., Duncan, A., Evans, T.D., Jones, K.R., Beyer, H.L., Schuster, R., Walston, J., Ray, J.C., Robinson, J.G., Callow, M., Clements, T., Costa, H.M., DeGemmis, A., Elsen, P.R., Ervin, J., Franco, P., Goldman, E., Goetz, S., Hansen, A., Hofsvang, E., Jantz, P., Jupiter, S., Kang, A., Langhammer, P., Laurance, W.F., Lieberman, S., Linkie, M., Malhi, Y., Maxwell, S., Mendez, M., Mittermeier, R., Murray, N.J., Possingham, H., Radachowsky, J., Saatchi, S., Samper, C., Silverman, J., Shapiro, A., Strassburg, B., Stevens, T., Stokes, E., Taylor, R., Tear, T., Tizard, R., Venter, O., Visconti, P., Wang, S., Watson, J.E.M. (2020) Anthropogenic modification of forests means only 40% of remaining forests have high ecosystem integrity. Nature Communications 11, 5978.

Gurney, G.G., Adams, V.M., Álvarez-Romero, J.G., Claudet, J. (2023) Area-based conservation: Taking stock and looking ahead. One Earth 6, 98–104.

Guizar-Coutiño, A., Jones, J.P.G., Balmford, A., Carmenta, R., Coomes, D.A. (2022) A global evaluation of the effectiveness of voluntary REDD+ projects at reducing deforestation and degradation in the moist tropics. Conservation Biology 36, e13970.

Hansen, M. C., Wang, L., Song, X.-P., Tyukavina, A., Turubanova, S., Potapov, P. V., & Stehman, S. V. The fate of tropical forest fragments. Science Advances, 6(11), eaax8574.

Harris, N. L., Gibbs, D. A., Baccini, A., Birdsey, R. A., de Bruin, S., Farina, M., … Tyukavina, A. (2021). Global maps of twenty-first century forest carbon fluxes. Nature Climate Change, 11(3), 234–240.

Herrmann, S.M., Brandt, M., Rasmussen, K., Fensholt, R. (2020) Accelerating land cover change in West Africa over four decades as population pressure increased. Communications Earth & Environment 1, 53.

Hoffman, M., Koenig, K., Bunting, G., Costanza, J., & Williams, K. J. (2016). Biodiversity Hotspots (version 2016.1). Zenodo.

Holloway V, and E Giandomenico. 2009. The History of REDD Policy, Carbon Planet White Paper, p. 20. Available at http://unfccc.int/files/methods_science/redd/application/pdf/the_history_of_redd_carbon_planet.pdf.

PBES. (2019). Global assessment report on biodiversity and ecosystem services of the Intergovernmental Science-Policy Platform on Biodiversity and Ecosystem Services (Version 1). Zenodo.

ayachandran, S., De Laat, J., Lambin, E.F., Stanton, C.Y., Audy, R., Thomas, N.E. (2017) Cash for carbon: A randomized trial of payments for ecosystem services to reduce deforestation. Science 357, 267–273.

oppa, L., & Pfaff, A. (2010). Reassessing the forest impacts of protection. Annals of the New York Academy of Sciences, 1185(1), 135–149.

KBA Partnership. Key Biodiversity Areas: Standards and Guidelines for Identifying KBAs. 2024. KBA Partnership, https://www.keybiodiversityareas.org/.

Kleinschroth, F., Laporte, N., Laurance, W.F., Goetz, S.J., Ghazoul, J. (2019) Road expansion and persistence in forests of the Congo Basin. Nature Sustainability 2, 628–634.

Laurance, W. F., Albernaz, A. K., Schroth, G., Fearnside, P. M., Bergen, S., Venticinque, E. M., & Da Costa, C. (2002). Predictors of deforestation in the Brazilian Amazon. Journal of Biogeography, 29(5-6), 737–748.

Laurance, W. F., Campbell, M. J., Alamgir, M., & Mahmoud, M. I. (2017). Road Expansion and the Fate of Africa’s Tropical Forests. Frontiers in Ecology and Evolution, 5.

Ma, J., Li, J., Wu, W., Liu, J. (2023) Global forest fragmentation change from 2000 to 2020. Nature Communications 14, 3752.

Malley, J. D., Kruppa, J., Dasgupta, A., Malley, K. G., & Ziegler, A. (2012). Probability Machines. Consistent Probability Estimation Using Nonparametric Learning Machines, 51(01), 74–81.

Maxwell, S. L., Evans, T., Watson, J. E. M., Morel, A., Grantham, H., Duncan, A., … Malhi, Y. (2019). Degradation and forgone removals increase the carbon impact of intact forest loss by 626%. Science Advances, 5(10), eaax2546.

Malan, M., Carmenta, R., Gsottbauer, E., Hofman, P., Kontoleon, A., Swinfield, T., Voors, M. (2024) Evaluating the impacts of a large-scale voluntary REDD+ project in Sierra Leone. Nature Sustainability.

Myers, N., Mittermeier, R.A., Mittermeier, C.G., da Fonseca, G.A.B., Kent, J. (2000) Biodiversity hotspots for conservation priorities. Nature 403, 853.

Pendrill, F., Gardner, T.A., Meyfroidt, P., Persson, U.M., Adams, J., Azevedo, T., Bastos Lima, M.G., Baumann, M., Curtis, P.G., De Sy, V., Garrett, R., Godar, J., Goldman, E.D., Hansen, M.C., Heilmayr, R., Herold, M., Kuemmerle, T., Lathuillière, M.J., Ribeiro, V., Tyukavina, A., Weisse, M.J., West, C. (2022) Disentangling the numbers behind agriculture-driven tropical deforestation. Science 377, eabm9267.

Pfaff, A. S. (1999). What drives deforestation in the Brazilian Amazon?: Evidence from satellite and socioeconomic data. Journal of environmental economics and management, 37(1), 26–43.

Pillay, R., Venter, M., Aragon-Osejo, J., González-del-Pliego, P., Hansen, A. J., Watson, J. E. M., & Venter, O. (2022). Tropical forests are home to over half of the world’s vertebrate species. Frontiers in Ecology and the Environment, 20(1), 10–15.

Plumptre, A.J., Baisero, D., Brooks, T.M., Buchanan, G., Butchart, S.H.M., Bowser, A., Boyd, C., Carneiro, A.P.B., Davies, T., Elliot, W., Foster, M., Langhammer, P.F., Marnewick, D., Matiku, P., McCreless, E., Raudsepp-Hearne, C., Tordoff, A.W., Azpiroz, A.B., Trisurat, Y., Upgren, A. (2024) Targeting site conservation to increase the effectiveness of new global biodiversity targets. One Earth 7, 11–17.

Powlen, K.A., Salerno, J., Jones, K.W., Gavin, M.C. (2023) Identifying socioeconomic and biophysical factors driving forest loss in protected areas. Conservation Biology 37, e14058.

Pya, N., & Wood, S. N. (2015). Shape constrained additive models. Statistics and Computing, 25(3), 543–559.

R Core Team (2023). R: A language and environment for statistical computing. R Foundation for Statistical Computing, Vienna, Austria. URL https://www.R-project.org/.

Simkins, A. T., Donald, P. F., Beresford, A. E., Butchart, S. H. M., Fa, J. E., Fernández-Llamazares, A. O., … Buchanan, G. M. (2024). Rates of tree cover loss in key biodiversity areas on Indigenous Peoples’ lands. Conservation Biology, 38(3), e14195.

Simonet, G., Subervie, J., Ezzine-de-Blas, D., Cromberg, M., Duchelle, A.E. (2019) Effectiveness of a REDD+ project in reducing deforestation in the Brazilian Amazon. American Journal of Agricultural Economics 101, 211–229.

Thompson, I. D., Guariguata, M. R., Okabe, K., Bahamondez, C., Nasi, R., Heymell, V., & Sabogal, C. (2013). An operational framework for defining and monitoring forest degradation. Ecology and Society, 18(2).

Trew, B. T., Edwards, D. P., Lees, A. C., Klinges, D. H., Early, R., Svátek, M., … Maclean, I. M. D. (2024). Novel temperatures are already widespread beneath the world’s tropical forest canopies. Nature Climate Change, 1–7.

Trew, B. T., Lees, A. C., Edwards, D. P., Early, R., & Maclean, I. M. D. (2024). Identifying climate-smart tropical Key Biodiversity Areas for protection in response to widespread temperature novelty. Conservation Letters.

Tracewski, Ł., Butchart, S. H., Di Marco, M., Ficetola, G. F., Rondinini, C., Symes, A., … Buchanan, G. M. (2016). Toward quantification of the impact of 21st-century deforestation on the extinction risk of terrestrial vertebrates. Conserv Biol, 30(5), 1070–1079.

Tyukavina, A., Hansen, M. C., Potapov, P. V., Krylov, A. M., & Goetz, S. J. (2016). Pan-tropical hinterland forests: mapping minimally disturbed forests. Global Ecology and Biogeography, 25(2), 151–163.

Weiss, D.J., Nelson, A., Gibson, H.S., Temperley, W., Peedell, S., Lieber, A., Hancher, M., Poyart, E., Belchior, S., Fullman, N., Mappin, B., Dalrymple, U., Rozier, J., Lucas, T.C.D., Howes, R.E., Tusting, L.S., Kang, S.Y., Cameron, E., Bisanzio, D., Battle, K.E., Bhatt, S., Gething, P.W. (2018) A global map of travel time to cities to assess inequalities in accessibility in 2015. Nature 553, 333–336.

Wright, M. N. & Ziegler, A. (2017). ranger: A fast implementation of random forests for high dimensional data in C++ and R. J Stat Softw 77:1–17.

Wolf, C., Levi, T., Ripple, W. J., Zárrate-Charry, D. A., & Betts, M. G. (2021). A forest loss report card for the world’s protected areas. Nature Ecology & Evolution, 5(4), 520–529.

Vancutsem, C., Achard, F., Pekel, J.F., Vieilledent, G., Carboni, S., Simonetti, D., Gallego, J., Aragão, L.E.O.C., Nasi, R. (2021) Long-term (1990–2019) monitoring of forest cover changes in the humid tropics. Science Advances 7, eabe1603.

Yates, L. A., Aandahl, Z., Richards, S. A., & Brook, B. W. (2023). Cross validation for model selection: A review with examples from ecology. Ecological Monographs, 93(1), e1557.

Zanella, L., Folkard, A. M., Blackburn, G. A., & Carvalho, L. M. T. (2017). How well does random forest analysis model deforestation and forest fragmentation in the Brazilian Atlantic forest? Environmental and Ecological Statistics, 24(4), 529–549.

