## Supplementary Information for "Predicting near-future deforestation in West African Key Biodiversity Areas to inform conservation urgency"

1.0 Supplementary Methods.	2

2.0 Supplementary Results.	5

3.0 Supplementary Tables.	6

4.0 Supplementary Figures.	25

### 1.0 Supplementary Methods.

**Natural features**

For each model group, a digital elevation model (DEM) at 60 m gridded resolution was downloaded from Amazon Web Services (available at: https://aws.amazon.com/public-datasets/terrain/) and converted to elevational bands at 100 m intervals. The DEM was then used to calculate mean slope horizontal to the ground (degrees) across the landscape using the *microclima* package for R (Maclean et al., 2019; R Core Team, 2023).

**Forest access**

For each model group, road density was calculated as the total length of road for each grid cell in the landscape within a 2 km^2^ moving window; a window size chosen to capture road density locality. Data on the locations of paved highways and tracks was obtained from Open Source Mapper (available at: https://www.openstreetmap.org). Additionally, to derive a measure of forest accessibility, we estimated the relative cost (as time taken) of a vehicle moving to each forest grid cell from the nearest settlement. Data on the locations of settlements - including villages, towns and cities - was obtained from Open Source Mapper. First, slope was calculated from altitude and distance between cell centres using the digital elevation model described above and the *gdistance* package for R (van Etten, 2017; R Core Team, 2023). A cost-surface layer dependent on slope was then derived based on the wheeled-vehicle critical slope cost function and a terrain multiplication factor (*sensu* Alberti, 2019). Additionally, a terrain multiplication factor was defined as the multiplier cost of a vehicle moving through different habitat or terrain and was derived from combining information on forest cover for the explanatory time period (Vancutsem et al., 2021) as well as road and river locations and other land cover. Information on river locations was obtained from the WWF Hydrosheds database (Lehner & Grill, 2013) and the location of other land cover types was obtained from the European Space Agency Land Cover data for 2020 (ESA, 2020). Forest, water, paved roads, tracks, paths and other land cover were assigned a terrain multiplication factor to represent the relative difference in cost of moving across different terrain. Finally, an accumulated cost surface was derived as the minimum least-cost distance from the nearest settlement. To derive forest access layers for future timesteps, cells which were deemed ‘deforested’ were converted to non forest for the terrain multiplication factor and the accumulated cost surface was calculated as above.

**Anthropic influences**

Neighbouring deforestation for each forest cell was calculated as the proportion of deforestation surrounding a focal cell within a 305 m moving window, i.e. the proportion of deforestation within 122 m of the grid cell, a distance chosen to represent close proximity. Deforestation was determined by summing deforestation events recorded in the explanatory time period using the deforestation year dataset from TMF JRC (Vancutsem et al., 2021). The same process was repeated for degradation events to derive spatial patterns of neighbouring degradation. To derive new layers for neighbouring deforestation and degradation at future timesteps, binary projections of deforestation and degradation events were used to calculate the metrics as above.

To assess the impact of subnational administrations on spatial patterns of deforestation, we calculated the average number of annual deforestation events over a ten-year baseline period (the ten years prior to the prediction time period) and over a two-year recent period (the two years prior to the prediction time period) for each subnational administration. Subnational administrative boundaries were freely obtained from the COD - Subnational Administrative Boundaries database (UNDP, 2023). The difference between the average annual deforestation events for the two-year and ten-year time periods was used as an anomaly estimate for the most recent two years. The same process was repeated to derive an anomaly estimate for annual degradation by subnational administrations. To derive new anomaly estimates for deforestation and degradation at future timesteps, binary projections of deforestation and degradation events were used to calculate the metrics as above.

**Conservation**

To derive an explanatory layer for Protected Area Designation, protected area boundaries were obtained from the World Database of Protected Areas (Protected Planet, 2021) and cleaned using standard protocols (Hanson, 2022). PA designations were classified 1 - 7 based on the IUCN categories (Ia & Ib = 1, II = 2, III = 3, IV = 4, V = 5, VI = 6 and None = 7).

**Forest landscape**

A range of landscape metrics were chosen as explanatory variables to represent forest fragmentation, connectivity and forest extent: (i) contiguity Index, (ii) radius of gyration, (iii) perimeter-area ratio, (iv) core area index, and (v) Euclidean nearest neighbour distance. Annual change data from JRC (Vancutsem et al., 2021) was converted to binary forest or non-forest maps (1/0), where both undisturbed and degraded forest were classed as forest. These forest maps were then processed as patch-type landscape metrics using the *landscapemetrics* package for R (Hesselbarth, 2019; R Core, 2023). We chose to include a selection of metrics which would represent all aspects of forest integrity, including forest connectivity, patch size and edge effects. To derive new landscape metrics at future timesteps, binary projections of deforestation were overlaid with the most recent forest/non forest map (above) and deforested cells converted to non-forest, with the resulting forest map used to calculate the metrics as above.

**Thresholding deforestation & degradation probability**

To derive a threshold for classifying the continuous output from the models into a binary output of whether a deforestation or degradation event is projected to occur, a historical sensitivity analysis was performed for each model group. Using the calibrated models, weighted ensemble mean probabilities of deforestation and degradation were produced for the years 2017, 2019 and 2021. The ensemble probabilities were classed as an occurrence of deforestation or degradation at multiple threshold values: from the 70th to 98th percentile values (at 2 percentile steps) of all the trained models. To minimise missed positive deforestation occurrences, we used percentiles as threshold values which accommodates each model group’s probability distribution as well as highly unbalanced datasets (i.e. few deforestation events compared to no events). The hindcast and thresholded projections of occurrences were then used to create confusion matrices (n = 6) corresponding to observed historical occurrences (n = 3). Each individual threshold matrix was then tested against performance metrics and an F1 score (balancing precision and recall) was derived for each threshold value (Kuhn, 2008; R Core Team, 2023).

The percentile values with the maximum F1 score across years were averaged across the testing years and used to define the occurrence of deforestation or degradation for the ensemble projections of deforestation and degradation probability for 2024 to 2025 (inclusive). These binary projections of events or non-events were then used to derive new forest cover maps for the start of 2026 and ergo new dynamic explanatory variables as per the methods. Projections of deforestation and degradation events for 2026 to 2027 (inclusive) were derived as above and this process was repeated every two years up to the end of 2033.

**References**

Alberti, G. (2019). movecost: An R package for calculating accumulated slope-dependent anisotropic cost-surfaces and least-cost paths. SoftwareX, 10, 100331.

European Space Agency Climate Change Initiative (ESA) - Land Cover (2022) Land Cover Product, Version 2.1. Available at: https://climate.esa.int/en/projects/land-cover/.

Hanson J.O. (2022) wdpar: Interface to the World Database on Protected Areas. Journal of Open-Source Software, 7: 4594.

Herzog, I. (2020). Spatial analysis based on cost functions. In Archaeological spatial analysis (pp. 333-358). Routledge.

Hesselbarth, M., Sciaini, M., With, K., Wiegand, K., & Nowosad, J. (2019). landscapemetrics : an open‐source R tool to calculate landscape metrics. Ecography, 42, 1648-1657.

Kuhn, M. (2008) Building Predictive Models in R Using the caret Package. Journal of Statistical Software 28, 1 - 26.

Lehner, B., Grill G. (2013). Global river hydrography and network routing: baseline data and new approaches to study the world’s large river systems. Hydrological Processes, 27(15): 2171–2186.

Maclean, I. M. D., Mosedale, J. R., & Bennie, J. J. (2019). Microclima: An r package for modelling meso- and microclimate. Methods in Ecology and Evolution, 10(2), 280-290.

Protected Planet: The World Database on Protected Areas (WDPA) (2021). Available at: https://www.protectedplanet.net

R Core, T. (2023). R: A language and environment for statistical computing. . Vienna: R Foundation for Statistical Computing.

van Etten, J. (2017). R Package gdistance: Distances and Routes on Geographical Grids. Journal of Statistical Software, 76(13), 1 - 21.

Vancutsem, C., Achard, F., Pekel, J. F., Vieilledent, G., Carboni, S., Simonetti, D., . . . Nasi, R. (2021). Long-term (1990–2019) monitoring of forest cover changes in the humid tropics. Science Advances, 7(10), eabe1603.

United Nations Development Programme (UNDP) GeoHub. (2023). COD - Subnational Administrative Boundaries database. Available at: https://geohub.data.undp.org/data/eacf362c7f81ace4de7298bdb71341fa.

### 2.0 Supplementary Results.

**Model Evaluation**

Deforestation models performed very well overall whereby 95% of model groups had mean AUC scores of greater than 0.7 during testing of deforestation models (supplementary Table S3 and Fig. S2). The best performing deforestation model groups (those with minimum AUC scores greater than 0.8) were predominantly large model groups with substantial forest extent, including a sizeable cluster of KBAs on the Liberia-Côte d'Ivoire border (model group 11), in southwestern Ghana (model group 15), across the Nigeria-Cameroon border (model group 22), in a largely forested region in Cameroon (model group 24) and across the Greater Gola Landscape in Liberia and Sierra Leone (model group 6). As expected, degradation was more difficult to predict. Nevertheless, 73% of model groups had mean AUC scores of greater than 0.7 during testing of degradation models.

### 3.0 Supplementary Tables.

**Table S1 |** Explanatory variables selected (indicated by X) for model building after stepwise VIF analysis.

| Model Group | Neighbouring deforestation | Neighbouring degradation | Mean slope | Elevation band | Motor access | Road density | Deforestation anomaly | ENN distance | Radius of gyration | Perimeter-area ratio | Core Area Index | Contiguity Index | Conservation features |
| --- | --- | --- | --- | --- | --- | --- | --- | --- | --- | --- | --- | --- | --- |
| Group 1 | x | x | x |  | x | x | x | x | x | x |  |  | x |
| Group 3 | x | x | x |  | x | x | x | x |  | x |  |  |  |
| Group 5 | x | x | x | x |  | x | x | x | x |  |  |  |  |
| Group 6 | x | x | x | x | x | x | x | x | x |  |  |  | x |
| Group 7 | x | x | x | x | x | x | x | x | x |  |  |  | x |
| Group 8 | x | x | x |  | x | x | x | x |  | x |  |  | x |
| Group 9 | x | x | x | x | x | x | x | x | x |  |  |  | x |
| Group 10 | x | x | x | x | x | x | x | x | x | x |  |  | x |
| Group 11 | x | x | x | x | x | x | x | x |  | x |  |  | x |
| Group 13 | x | x | x | x | x | x | x | x |  | x |  |  | x |
| Group 14 | x | x | x | x | x | x | x | x |  | x |  |  | x |
| Group 15 | x | x | x | x | x | x | x | x | x |  | x |  | x |
| Group 16 | x | x | x | x | x | x | x | x | x |  |  |  | x |
| Group 17 | x | x | x | x | x | x | x | x | x |  |  |  | x |
| Group 20 | x | x | x | x | x | x | x | x | x |  |  |  | x |
| Group 21 | x | x | x |  | x | x | x | x | x | x |  |  | x |
| Group 22 | x | x | x | x | x | x | x | x | x | x |  |  | x |
| Group 23 | x | x | x | x | x | x | x | x | x |  |  |  | x |
| Group 24 | x | x | x | x | x | x | x | x | x |  |  |  | x |
| Group 25 | x | x | x | x | x | x | x | x | x |  |  |  | x |
| Group 26 | x | x | x | x | x | x |  | x | x |  |  |  | x |
| Group 30 | x | x | x | x | x | x | x | x |  | x |  |  | x |

**Table S2 |** Minimum and maximum linear correlation coefficient between predictor variables selected for model building after stepwise VIF analysis.

|  | **Minimum Correlation** | | **Maximum Correlation** | |
| --- | --- | --- | --- | --- |
| **Group** | **Coefficient** | **Variables** | **Coefficient** | **Variables** |
| Group 1 | -0.003 | ENN distance ~ Neighbouring deforestation | 0.595 | Conservation features ~ Motor access |
| Group 3 | 0.001 | ENN distance ~ Neighbouring deforestation | 0.588 | Conservation feature ~ Motor access |
| Group 5 | -0.011 | Deforestation anomaly ~ ENN distance | 0.582 | Conservation feature ~ Motor access |
| Group 6 | -0.007 | ENN distance ~ Neighbouring deforestation | 0.613 | Conservation feature ~ Motor access |
| Group 7 | -0.002 | ENN distance ~ Elevation band | 0.588 | Conservation feature ~ Motor access |
| Group 8 | 0.002 | Deforestation anomaly ~ Road density | 0.602 | Conservation feature ~ Motor access |
| Group 9 | 0.000 | Deforestation anomaly ~ Road density | 0.600 | Conservation feature ~ Motor access |
| Group 10 | 0.001 | Mean slope ~ Elevation band | 0.571 | Conservation feature ~ Motor access |
| Group 11 | -0.004 | Deforestation anomaly ~ Road density | 0.597 | Conservation feature ~ Motor access |
| Group 13 | 0.006 | ENN distance ~ Neighbouring deforestation | 0.598 | Conservation feature ~ Motor access |
| Group 14 | -0.003 | ENN distance ~ Neighbouring deforestation | 0.577 | Conservation feature ~ Motor access |
| Group 15 | -0.001 | Neighbouring deforestation ~ Elevation band | 0.587 | Conservation feature ~ Motor access |
| Group 16 | 0.004 | ENN distance ~ Neighbouring deforestation | 0.571 | Conservation feature ~ Motor access |
| Group 17 | -0.002 | ENN distance ~ Elevation band | 0.586 | Conservation feature ~ Motor access |
| Group 20 | 0.000 | ENN distance ~ Elevation band | 0.606 | Conservation feature ~ Motor access |
| Group 21 | 0.000 | Deforestation anomaly ~ Road density | 0.576 | Conservation feature ~ Motor access |
| Group 22 | 0.000 | ENN distance ~ Elevation band | 0.603 | Conservation feature ~ Motor access |
| Group 23 | 0.000 | ENN distance ~ Neighbouring deforestation | 0.592 | Conservation feature ~ Motor access |
| Group 24 | 0.000 | Deforestation anomaly ~ Road density | 0.574 | Conservation feature ~ Motor access |
| Group 25 | 0.004 | Deforestation anomaly ~ Road density | 0.583 | Conservation feature ~ Motor access |
| Group 26 | 0.002 | ENN distance ~ Neighbouring deforestation | 0.592 | Conservation feature ~ Motor access |
| Group 30 | 0.001 | ENN distance ~ Neighbouring deforestation | 0.584 | Conservation feature ~ Motor access |

**Table S3 |** Summary statistics of the area under the curve (AUC) scores obtained for each model group during model testing for deforestation.

|  | **Deforestation Models** | | | | **Degradation Models** | | | | |
| --- | --- | --- | --- | --- | --- | --- | --- | --- | --- |
| **Model Group** | **Mean** | **Max.** | **Min.** | **SD** | | **Mean** | **Max.** | **Min.** | **SD** |
| group 1 | 0.79 | 0.9 | 0.7 | 0.06 | | 0.74 | 0.8 | 0.62 | 0.06 |
| group 10 | 0.71 | 0.73 | 0.7 | 0.01 | | 0.68 | 0.7 | 0.67 | 0.01 |
| group 11 | 0.85 | 0.87 | 0.84 | 0.01 | | 0.78 | 0.8 | 0.77 | 0.01 |
| group 13 | 0.79 | 0.83 | 0.75 | 0.03 | | 0.76 | 0.77 | 0.73 | 0.01 |
| group 14 | 0.76 | 0.78 | 0.74 | 0.01 | | 0.71 | 0.72 | 0.7 | 0.01 |
| group 15 | 0.81 | 0.82 | 0.8 | 0.01 | | 0.74 | 0.76 | 0.73 | 0.01 |
| group 16 | 0.77 | 0.8 | 0.72 | 0.02 | | 0.75 | 0.77 | 0.74 | 0.01 |
| group 17 | 0.72 | 0.75 | 0.68 | 0.02 | | 0.67 | 0.7 | 0.66 | 0.02 |
| group 20 | 0.8 | 0.82 | 0.78 | 0.01 | | 0.72 | 0.76 | 0.71 | 0.01 |
| group 21 | 0.78 | 0.84 | 0.73 | 0.03 | | 0.67 | 0.72 | 0.59 | 0.04 |
| group 22 | 0.88 | 0.89 | 0.86 | 0.01 | | 0.86 | 0.88 | 0.82 | 0.02 |
| group 23 | 0.8 | 0.82 | 0.76 | 0.02 | | 0.76 | 0.8 | 0.73 | 0.03 |
| group 24 | 0.86 | 0.88 | 0.85 | 0.01 | | 0.8 | 0.82 | 0.78 | 0.01 |
| group 25 | 0.78 | 0.83 | 0.7 | 0.04 | | 0.7 | 0.76 | 0.63 | 0.04 |
| group 26 | 0.75 | 0.78 | 0.71 | 0.02 | | 0.69 | 0.73 | 0.66 | 0.02 |
| group 3 | 0.75 | 0.81 | 0.71 | 0.04 | | 0.74 | 0.8 | 0.66 | 0.04 |
| group 30 | 0.92 | 0.97 | 0.85 | 0.04 | | 0.9 | 0.95 | 0.84 | 0.04 |
| group 5 | 0.68 | 0.7 | 0.66 | 0.01 | | 0.65 | 0.68 | 0.63 | 0.01 |
| group 6 | 0.81 | 0.83 | 0.8 | 0.01 | | 0.77 | 0.78 | 0.76 | 0.01 |
| group 7 | 0.8 | 0.82 | 0.78 | 0.01 | | 0.7 | 0.72 | 0.69 | 0.01 |
| group 8 | 0.72 | 0.77 | 0.68 | 0.03 | | 0.64 | 0.69 | 0.6 | 0.03 |
| group 9 | 0.81 | 0.84 | 0.77 | 0.02 | | 0.76 | 0.78 | 0.73 | 0.02 |

**Table S4 |** Mean importance of each explanatory variable across all model blocks (n = 10) and model groups (n = 22). Importance is defined as the increase in node purity. Importance values are an estimate of size of influence but do not indicate the direction of influence.

| **Predictor** | **Mean** | **SD** | **Sample size** |
| --- | --- | --- | --- |
| Neighbouring Deforestation | 1.0262 | 0.0130 | 220 |
| Conservation Feature | 1.0175 | 0.0311 | 200 |
| Radius of Gyration | 1.0152 | 0.0207 | 160 |
| Neighbouring Degradation | 1.0126 | 0.0124 | 220 |
| Motor Access | 1.0113 | 0.0181 | 210 |
| Core Area Index | 1.0097 | NA | 10 |
| Anomaly in Deforestation Rate | 1.0096 | 0.0184 | 210 |
| Perimeter-area Ratio | 1.0069 | 0.0088 | 100 |
| Road Density | 1.0028 | 0.0105 | 220 |
| Euclidean Nearest Neighbour Distance | 1.0000 | 0.0026 | 220 |
| Forest Contiguity | 1.0000 | NA | 10 |
| Elevation Band | 0.9988 | 0.0142 | 180 |
| Mean Slope | 0.9968 | 0.0067 | 220 |

**Table S5 |** Historical forest loss in Key Biodiversity Areas within each country across the Guinean Forests of West Africa between 2013 and 2023. Presented by country as total forest loss (ha) within all KBAs in each country and mean loss across all KBAs in each country. KBAs in Togo and Benin are not included in this study as model groups here did not perform sufficiently.

| **Country** | **Total Loss (Ha)** | **Total Loss (%)** | **KBA Loss (Ha)** | **KBA Loss (%)** |
| --- | --- | --- | --- | --- |
| Côte d’Ivoire | -1,524,566 | 33.91 | -121,791 | 14 |
| Nigeria | -934,873 | 15.2 | -32,981 | 5.07 |
| Ghana | -914,570 | 25.38 | -32,326 | 6.94 |
| Cameroon | -716,617 | 3.61 | -5,857 | 0.72 |
| Liberia | -640,565 | 7.8 | -44,289 | 1.64 |
| Sierra Leone | -518,034 | 28.99 | -18,291 | 10.68 |
| Guinea | -196,995 | 23.62 | -9,966 | 5.83 |
| Equatorial Guinea | -31,196 | 1.22 | -143 | 0.18 |

**Table S6 |** Forest loss within each KBA across the Guinean Forests of West Africa between 2013 and 2023 (inclusive).

| **KBA** | **Country** | **Mean Annual Loss (Ha)** | **Mean Annual Loss (%)** | **Total Loss (Ha)** | **Total Loss (%)** |
| --- | --- | --- | --- | --- | --- |
| Cavally and Goin - Debe Forest Reserves | Côte d'Ivoire | -7439 | 6.41 | -81827 | 52.48 |
| Omo Forest Reserve | Nigeria | -1655 | 1.74 | -18209 | 17.64 |
| Tano-Offin Forest Reserve | Ghana | -1113 | 3.07 | -12246 | 31.35 |
| Zwedru | Liberia | -1080 | 1.78 | -11884 | 18.52 |
| Gueoule and Glo Mountain Forest Reserves | Côte d'Ivoire | -1029 | 9.88 | -11315 | 69.54 |
| Kangari Hills Non-hunting Forest Reserve | Sierra Leone | -893 | 2.93 | -9823 | 28.06 |
| Peko Mountain National Park | Côte d'Ivoire | -792 | 8.23 | -8712 | 62.59 |
| Wologizi mountains | Liberia | -728 | 0.46 | -8008 | 4.9 |
| Mount Nimba Strict Nature Reserve | Côte d'Ivoire | -705 | 4.34 | -7753 | 38.85 |
| Okomu National Park | Nigeria | -530 | 0.67 | -5825 | 7.14 |
| Subri River Forest Reserve | Ghana | -514 | 0.97 | -5655 | 10.32 |
| Lofa-Gola-Mano Complex | Liberia | -373 | 0.09 | -4108 | 0.95 |
| Massif du Ziama | Guinea | -337 | 0.44 | -3703 | 4.7 |
| Cestos - Senkwen | Liberia | -322 | 0.09 | -3547 | 1.04 |
| Mabi Forest reserve | Côte d'Ivoire | -310 | 0.54 | -3408 | 5.74 |
| Gio National Forest | Liberia | -299 | 0.65 | -3291 | 6.91 |
| Foret Classe de Mont Bero | Guinea | -288 | 3.24 | -3166 | 30.74 |
| Western Area Peninsula Forest National Park | Sierra Leone | -278 | 1.87 | -3060 | 18.77 |
| Kpelle Forest | Liberia | -268 | 0.13 | -2945 | 1.37 |
| Mopri Forest Reserve | Côte d'Ivoire | -263 | 3.02 | -2891 | 29.19 |
| Wonegizi mountains | Liberia | -258 | 1 | -2835 | 10.43 |
| Tano-Ehuro Forest Reserve | Ghana | -252 | 3.26 | -2774 | 30.87 |
| Cross River National Park (Oban Division) | Nigeria | -237 | 0.09 | -2609 | 0.98 |
| Parc National de Taï et Réserve de faune du N'Zo | Côte d'Ivoire | -216 | 0.04 | -2378 | 0.45 |
| Bossematie Forest Reserve | Côte d'Ivoire | -197 | 0.97 | -2164 | 10.21 |
| Gashaka-Gumti National Park | Nigeria | -195 | 0.42 | -2146 | 4.55 |
| Loma Mountains Non-hunting Forest Reserve | Sierra Leone | -185 | 1.24 | -2039 | 12.89 |
| Konkouré | Guinea | -173 | 1.33 | -1906 | 13.71 |
| Cestos-Sapo South Corridor forest block | Liberia | -160 | 0.51 | -1755 | 5.48 |
| Afi River Forest Reserve | Nigeria | -158 | 0.34 | -1743 | 3.72 |
| Sapo - Grebo Corridor | Liberia | -154 | 0.08 | -1694 | 0.86 |
| Tchabal-Mbabo | Cameroon | -153 | 0.55 | -1684 | 5.93 |
| Grebo | Liberia | -130 | 0.05 | -1426 | 0.51 |
| Tingi Hills Non-hunting Forest Reserve | Sierra Leone | -126 | 3.86 | -1384 | 35.47 |
| Kambui Hills Forest Reserve | Sierra Leone | -119 | 0.54 | -1308 | 5.82 |
| Diécké | Guinea | -118 | 0.21 | -1302 | 2.32 |
| Eastern Bamenda highlands and associated hydrobasin | Cameroon | -118 | 0.46 | -1294 | 4.96 |
| Weeni creek and associated hydrobasin | Liberia | -112 | 1.15 | -1228 | 12 |
| Amansuri wetland | Ghana | -111 | 0.63 | -1225 | 6.74 |
| Neung South | Ghana | -109 | 1 | -1196 | 10.51 |
| Cross River National Park (Okwangwo Division) and Mbe Mountains | Nigeria | -108 | 0.13 | -1191 | 1.46 |
| Southern Scarp | Ghana | -99 | 1.12 | -1086 | 11.75 |
| West Nimba | Liberia | -92 | 0.91 | -1007 | 9.53 |
| Draw River Forest Reserve | Ghana | -89 | 0.52 | -978 | 5.54 |
| Krahn Bassa South | Liberia | -86 | 0.04 | -943 | 0.47 |
| Yapo and Mambo Forest Reserves | Côte d'Ivoire | -83 | 0.29 | -915 | 3.18 |
| Cestos Gbi | Liberia | -70 | 0.02 | -773 | 0.24 |
| Gola Forests | Sierra Leone | -70 | 0.09 | -769 | 1.03 |
| Atewa Range Forest Reserve | Ghana | -70 | 0.31 | -766 | 3.35 |
| Biseni forests | Nigeria | -69 | 0.33 | -755 | 3.62 |
| Bosomtwe Range Forest Reserve | Ghana | -67 | 0.98 | -741 | 10.37 |
| Kakum National Park - Assin Attandaso Resource Reserve | Ghana | -65 | 0.21 | -716 | 2.28 |
| Mount Cameroon and Mokoko-Onge | Cameroon | -60 | 0.07 | -663 | 0.75 |
| Pic de Fon | Guinea | -59 | 0.83 | -649 | 8.72 |
| Nimba mountains | Liberia | -55 | 0.49 | -609 | 5.31 |
| Ankasa Resource Reserve - Nini-Sushien National Park | Ghana | -55 | 0.12 | -605 | 1.31 |
| Yabassi | Cameroon | -50 | 0.02 | -552 | 0.21 |
| Cape Three Points Forest Reserve | Ghana | -49 | 1.75 | -544 | 17.66 |
| Bura River Forest Reserve | Ghana | -47 | 0.51 | -522 | 5.45 |
| Sierra Leone River Estuary | Sierra Leone | -46 | 0.52 | -503 | 5.58 |
| Fazao-Malfakassa National Park | Togo | -45 | 1.22 | -492 | 12.7 |
| Bakossi mountains | Cameroon | -45 | 0.06 | -490 | 0.65 |
| Boin River Forest Reserve | Ghana | -40 | 0.14 | -435 | 1.49 |
| Cestos-Sapo North Corridor forest blocks | Liberia | -38 | 0.05 | -420 | 0.52 |
| Tano-Nimiri Forest Reserve | Ghana | -36 | 0.19 | -393 | 2.1 |
| Bia National Park and Resource Reserve | Ghana | -34 | 0.11 | -374 | 1.2 |
| Santchou Faunal Reserve | Cameroon | -34 | 0.63 | -370 | 6.68 |
| Tano-Anwia Forest Reserve | Ghana | -31 | 0.23 | -345 | 2.53 |
| Yoyo River Forest Reserve | Ghana | -31 | 0.15 | -337 | 1.69 |
| Upper Orashi forests | Nigeria | -30 | 0.31 | -333 | 3.38 |
| Fure River Forest Reserve | Ghana | -30 | 0.22 | -330 | 2.39 |
| Jema-Asemkrom Forest Reserve | Ghana | -30 | 0.5 | -327 | 5.42 |
| Monts Nimba (part of Mount Nimba transboundary AZE) | Guinea | -25 | 0.23 | -279 | 2.51 |
| Mamiri Forest Reserve | Ghana | -24 | 0.53 | -262 | 5.66 |
| Mount Oku | Cameroon | -21 | 0.59 | -231 | 6.32 |
| Obudu Plateau | Nigeria | -19 | 2.48 | -210 | 24.28 |
| Boin Tano Forest Reserve | Ghana | -17 | 0.15 | -190 | 1.59 |
| Sangbe Mountain National Park | Côte d'Ivoire | -15 | 2.54 | -170 | 25.12 |
| Dadieso Forest Reserve | Ghana | -15 | 0.11 | -164 | 1.17 |
| Ebi River Shelterbelt Forest Reserve | Ghana | -14 | 1.07 | -152 | 11.21 |
| Basilé Peak National Park | Equatorial Guinea | -13 | 0.02 | -147 | 0.18 |
| Mont Nlonako | Cameroon | -13 | 0.02 | -146 | 0.23 |
| Grand Kru SouthWest blocks | Liberia | -13 | 0.02 | -138 | 0.25 |
| Kabitaï | Guinea | -12 | 2.54 | -128 | 25.13 |
| Mount Afadjato - Agumatsa Range forest | Ghana | -11 | 1.86 | -119 | 18.91 |
| Banyang Mbo Wildlife Sanctuary | Cameroon | -10 | 0.02 | -115 | 0.17 |
| Pra-Sushien Forest Reserve | Ghana | -10 | 0.05 | -112 | 0.6 |
| Yawri Bay | Sierra Leone | -10 | 1.07 | -111 | 11.21 |
| Grand Kru SouthEast Forest blocks | Liberia | -10 | 0.01 | -107 | 0.12 |
| Korup National Park | Cameroon | -9 | 0.01 | -95 | 0.07 |
| Azagny National Park | Côte d'Ivoire | -9 | 0.06 | -94 | 0.62 |
| Tanoe Forest Swamp Forest | Côte d'Ivoire | -7 | 0.06 | -81 | 0.68 |
| Luba Caldera Scientific Reserve | Equatorial Guinea | -7 | 0.01 | -79 | 0.16 |
| Adiopodoume | Côte d'Ivoire | -6 | 3.17 | -63 | 30.2 |
| Mont Manengouba | Cameroon | -5 | 0.13 | -59 | 1.45 |
| Mount Mbam | Cameroon | -5 | 0.67 | -50 | 7.13 |
| Mbi Crater Faunal Reserve - Mbingo forest | Cameroon | -4 | 0.78 | -39 | 8.31 |
| Misahöhe Forest Reserve | Togo | -3 | 1.16 | -28 | 12.09 |
| Mount Rata and Rumpi Hills Forest Reserve | Cameroon | -2 | 0.01 | -26 | 0.06 |
| Bali-Ngemba Forest Reserve | Cameroon | -2 | 0.52 | -24 | 5.57 |
| Bamboutos Mountains | Cameroon | -2 | 0.31 | -21 | 3.41 |
| Ngel-Nyaki Forest Reserve | Nigeria | -2 | 20.39 | -21 | 99.58 |
| Lamto Ecological Research Station | Côte d'Ivoire | -2 | 0.65 | -18 | 6.92 |
| Chutes de la Sala | Guinea | -1 | 2.21 | -16 | 22.6 |
| Banco National Park | Côte d'Ivoire | -1 | 0.04 | -13 | 0.43 |
| Njinsing - Tabenken | Cameroon | -1 | 0.47 | -13 | 5.03 |
| Nsuensa Forest Reserve | Ghana | -1 | 0.02 | -11 | 0.18 |
| Lake Nokoué | Benin | -1 | 0.47 | -7 | 5.07 |
| Sapo | Liberia | -1 | 0 | -7 | 0 |
| Mount Lefo | Cameroon | -1 | 2.75 | -6 | 26.72 |
| Mount Kupe | Cameroon | 0 | 0.01 | -1 | 0.12 |
| Mont Bana | Cameroon | 0 | 0.04 | 0 | 0.47 |
| Sapawsu Forest Reserve | Ghana | 0 | 3.62 | 0 | 36.36 |

**Table S7 |** Predicted mean forest loss in Key Biodiversity Areas within each country across the Guinean Forests of West Africa between 2025 and 2033 (inclusive). KBAs in Togo and Benin are not included in this study as model groups here did not perform sufficiently.

| **Country** | **Mean Predicted Forest Loss (Ha)** | **Mean Predicted Forest Loss (%)** |
| --- | --- | --- |
| Liberia | -6390 | 9.73 |
| Côte d'Ivoire | -2308 | 9.55 |
| Cameroon | -599 | 6.52 |
| Nigeria | -5134 | 5.24 |
| Ghana | -167 | 1.38 |
| Guinea | -150 | 1.2 |
| Equatorial Guinea | -390 | 0.64 |

**Table S8 |** Predicted forest loss within each KBA across the Guinean Forests of West Africa between 2025 and 2033 (inclusive).

| **KBA** | **Country** | **Mean Annual Forest Loss (Ha)** | **Mean Annual Forest Loss (%)** | **Total Loss (Ha)** | **Total Loss (%)** |
| --- | --- | --- | --- | --- | --- |
| Zwedru | Liberia | -592 | -1.53 | -48323 | 92.1 |
| Cross River National Park (Oban Division) | Nigeria | -969 | 0.37 | -24548 | 9.28 |
| Kpelle Forest | Liberia | -829 | 0.39 | -15154 | 7.14 |
| Lofa-Gola-Mano Complex | Liberia | -779 | 0.18 | -14991 | 3.49 |
| Wologizi mountains | Liberia | -573 | 0.37 | -12633 | 8.14 |
| Bossematie Forest Reserve | Côte d'Ivoire | -351 | 1.74 | -12053 | 63.24 |
| Gio National Forest | Liberia | -531 | 1.22 | -11493 | 25.84 |
| Cavally and Goin - Debe Forest Reserves | Côte d'Ivoire | -93 | -0.02 | -10304 | 13.87 |
| Mabi Forest reserve | Côte d'Ivoire | -651 | 1.19 | -10230 | 18.22 |
| Cestos Gbi | Liberia | -469 | 0.15 | -9828 | 3.11 |
| Omo Forest Reserve | Nigeria | -272 | 0.32 | -5950 | 6.99 |
| Cross River National Park (Okwangwo Division) and Mbe Mountains | Nigeria | -208 | 0.26 | -5510 | 6.85 |
| Gashaka-Gumti National Park | Nigeria | -201 | 0.45 | -5385 | 11.93 |
| Western Area Peninsula Forest National Park | Sierra Leone | -173 | 1.31 | -4356 | 32.89 |
| Afi River Forest Reserve | Nigeria | -113 | 0.24 | -4065 | 8.99 |
| Kangari Hills Non-hunting Forest Reserve | Sierra Leone | -180 | 0.72 | -3728 | 14.81 |
| West Nimba | Liberia | -157 | 1.65 | -3341 | 34.78 |
| Sapo - Grebo Corridor | Liberia | -216 | 0.11 | -3141 | 1.61 |
| Yabassi | Cameroon | -144 | 0.05 | -2515 | 0.95 |
| Tchabal-Mbabo | Cameroon | -104 | 0.39 | -2329 | 8.72 |
| Bakossi mountains | Cameroon | -85 | 0.11 | -2153 | 2.87 |
| Tano-Offin Forest Reserve | Ghana | -64 | 0.24 | -1620 | 6.04 |
| Korup National Park | Cameroon | -65 | 0.05 | -1092 | 0.84 |
| Yapo and Mambo Forest Reserves | Côte d'Ivoire | -171 | 0.61 | -1048 | 3.74 |
| Eastern Bamenda highlands and associated hydrobasin | Cameroon | -38 | 0.15 | -1011 | 4.05 |
| Wonegizi mountains | Liberia | -29 | 0.12 | -993 | 4.08 |
| Gola Forests | Sierra Leone | -32 | 0.04 | -886 | 1.2 |
| Amansuri wetland | Ghana | -35 | 0.21 | -765 | 4.49 |
| Konkouré | Guinea | -36 | 0.3 | -742 | 6.18 |
| Mont Nlonako | Cameroon | -42 | 0.07 | -607 | 0.95 |
| Okomu National Park | Nigeria | -28 | 0.04 | -493 | 0.65 |
| Grebo | Liberia | -43 | 0.02 | -473 | 0.17 |
| Bosomtwe Range Forest Reserve | Ghana | -16 | 0.24 | -455 | 7.03 |
| Atewa Range Forest Reserve | Ghana | -18 | 0.08 | -439 | 1.97 |
| Mount Cameroon and Mokoko-Onge | Cameroon | -28 | 0.03 | -428 | 0.48 |
| Basilé Peak National Park | Equatorial Guinea | -25 | 0.03 | -392 | 0.48 |
| Luba Caldera Scientific Reserve | Equatorial Guinea | -25 | 0.05 | -389 | 0.79 |
| Nimba mountains | Liberia | -7 | 0.06 | -357 | 3.27 |
| Sierra Leone River Estuary | Sierra Leone | -16 | 0.19 | -350 | 4.07 |
| Cestos-Sapo North Corridor forest blocks | Liberia | -27 | 0.03 | -348 | 0.43 |
| Mount Nimba Strict Nature Reserve | Côte d'Ivoire | -13 | 0.11 | -347 | 2.84 |
| Parc National de Taï et Réserve de faune du N'Zo | Côte d'Ivoire | -7 | 0 | -339 | 0.07 |
| Banyang Mbo Wildlife Sanctuary | Cameroon | -21 | 0.03 | -313 | 0.45 |
| Diécké | Guinea | -24 | 0.04 | -259 | 0.47 |
| Krahn Bassa South | Liberia | -14 | 0.01 | -257 | 0.13 |
| Santchou Faunal Reserve | Cameroon | -1 | 0.02 | -214 | 4.12 |
| Mbi Crater Faunal Reserve - Mbingo forest | Cameroon | -13 | 3.19 | -209 | 47.5 |
| Upper Orashi forests | Nigeria | -3 | 0.03 | -208 | 2.16 |
| Mopri Forest Reserve | Côte d'Ivoire | -7 | 0.09 | -189 | 2.69 |
| Yawri Bay | Sierra Leone | -2 | 0.18 | -181 | 20.51 |
| Mount Rata and Rumpi Hills Forest Reserve | Cameroon | -11 | 0.02 | -178 | 0.39 |
| Jema-Asemkrom Forest Reserve | Ghana | -5 | 0.08 | -168 | 2.92 |
| Bura River Forest Reserve | Ghana | -1 | 0.01 | -161 | 1.76 |
| Mount Oku | Cameroon | -7 | 0.21 | -155 | 4.5 |
| Tano-Anwia Forest Reserve | Ghana | -6 | 0.04 | -152 | 1.14 |
| Mamiri Forest Reserve | Ghana | -1 | 0.01 | -143 | 3.23 |
| Kambui Hills Forest Reserve | Sierra Leone | 0 | 0 | -136 | 0.64 |
| Tano-Nimiri Forest Reserve | Ghana | -1 | 0.01 | -132 | 0.72 |
| Foret Classe de Mont Bero | Guinea | -8 | 0.11 | -115 | 1.61 |
| Lamto Ecological Research Station | Côte d'Ivoire | -2 | 0.64 | -97 | 38.33 |
| Mount Mbam | Cameroon | -5 | 0.77 | -79 | 12.1 |
| Cape Three Points Forest Reserve | Ghana | -3 | 0.12 | -79 | 3.09 |
| Ankasa Resource Reserve - Nini-Sushien National Park | Ghana | -9 | 0.02 | -76 | 0.17 |
| Bali-Ngemba Forest Reserve | Cameroon | -5 | 1.16 | -73 | 17.71 |
| Pic de Fon | Guinea | -2 | 0.03 | -70 | 1.03 |
| Neung South | Ghana | -2 | 0.02 | -52 | 0.5 |
| Weeni creek and associated hydrobasin | Liberia | -1 | 0.02 | -47 | 0.51 |
| Biseni forests | Nigeria | -2 | 0.01 | -44 | 0.21 |
| Southern Scarp | Ghana | -2 | 0.02 | -44 | 0.53 |
| Tano-Ehuro Forest Reserve | Ghana | -2 | 0.03 | -36 | 0.57 |
| Boin River Forest Reserve | Ghana | -2 | 0.01 | -31 | 0.11 |
| Ebi River Shelterbelt Forest Reserve | Ghana | -4 | 0.32 | -29 | 2.34 |
| Boin Tano Forest Reserve | Ghana | -1 | 0.01 | -26 | 0.22 |
| Draw River Forest Reserve | Ghana | -2 | 0.01 | -21 | 0.13 |
| Fure River Forest Reserve | Ghana | -3 | 0.02 | -20 | 0.15 |
| Subri River Forest Reserve | Ghana | 0 | 0 | -19 | 0.04 |
| Dadieso Forest Reserve | Ghana | 0 | 0 | -18 | 0.13 |
| Cestos - Senkwen | Liberia | -1 | 0 | -16 | 0 |
| Loma Mountains Non-hunting Forest Reserve | Sierra Leone | 0 | 0 | -16 | 0.12 |
| Yoyo River Forest Reserve | Ghana | -2 | 0.01 | -11 | 0.06 |
| Gueoule and Glo Mountain Forest Reserves | Côte d'Ivoire | 0 | 0.01 | -11 | 0.22 |
| Bamboutos Mountains | Cameroon | -1 | 0.12 | -10 | 1.57 |
| Monts Nimba (part of Mount Nimba transboundary AZE) | Guinea | 0 | 0 | -10 | 0.1 |
| Kakum National Park - Assin Attandaso Resource Reserve | Ghana | -1 | 0 | -9 | 0.03 |
| Cestos-Sapo South Corridor forest block | Liberia | 0 | 0 | -7 | 0.02 |
| Tingi Hills Non-hunting Forest Reserve | Sierra Leone | 0 | 0 | -7 | 0.28 |
| Tanoe Forest Swamp Forest | Côte d'Ivoire | 0 | 0 | -5 | 0.04 |
| Mont Manengouba | Cameroon | 0 | 0 | -3 | 0.06 |
| Mount Lefo | Cameroon | 0 | -0.16 | -3 | 16.67 |
| Kabitaï | Guinea | 0 | 0.04 | -1 | 0.19 |
| Obudu Plateau | Nigeria | 0 | 0 | -1 | 0.11 |
| Adiopodoume | Côte d'Ivoire | 0 | 0 | 0 | 0 |
| Azagny National Park | Côte d'Ivoire | 0 | 0 | 0 | 0 |
| Banco National Park | Côte d'Ivoire | 0 | 0 | 0 | 0 |
| Bia National Park and Resource Reserve | Ghana | 0 | 0 | 0 | 0 |
| Chutes de la Sala | Guinea | 0 | 0 | 0 | 0 |
| Grand Kru SouthEast Forest blocks | Liberia | 0 | 0 | 0 | 0 |
| Grand Kru SouthWest blocks | Liberia | 0 | 0 | 0 | 0 |
| Massif du Ziama | Guinea | 0 | 0 | 0 | 0 |
| Mont Bana | Cameroon | 0 | 0 | 0 | 0 |
| Mount Kupe | Cameroon | 0 | 0 | 0 | 0 |
| Nsuensa Forest Reserve | Ghana | 0 | 0 | 0 | 0 |
| Peko Mountain National Park | Côte d'Ivoire | 0 | 0 | 0 | 0 |
| Pra-Sushien Forest Reserve | Ghana | 0 | 0 | 0 | 0 |
| Sangbe Mountain National Park | Côte d'Ivoire | 0 | 0 | 0 | 0 |
| Sapawsu Forest Reserve | Ghana | 0 | 0 | 0 | 0 |
| Sapo | Liberia | 0 | 0 | 0 | 0 |

### 4.0 Supplementary Figures.

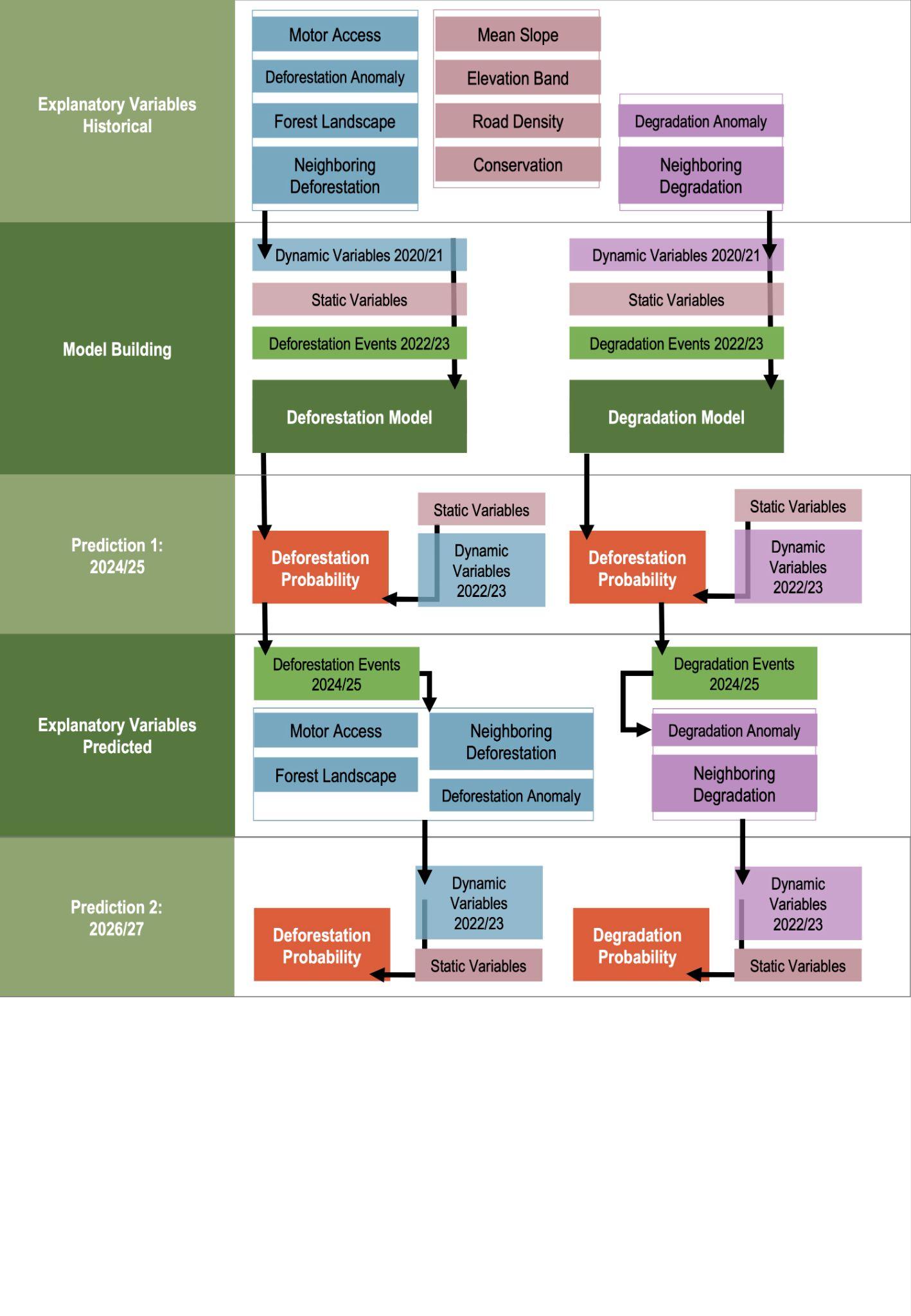

**Figure S1 |** Full workflow for modelling near-future deforestation risk using a stepwise machine-learning method. Dynamic explanatory variables associated with deforestation are represented in blue and those associated with degradation in purple. Static variables are represented in pink.

**A**
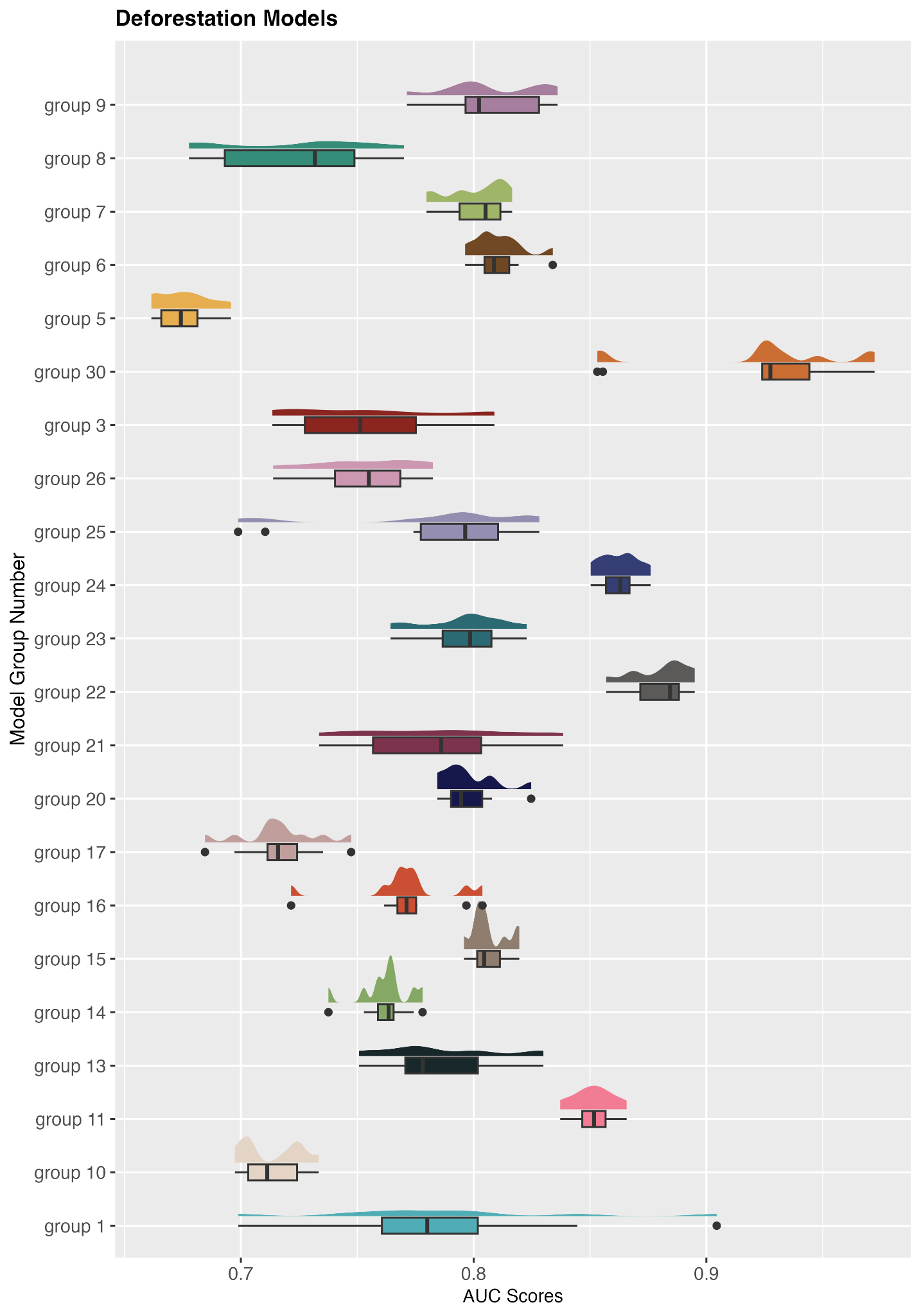
**B**
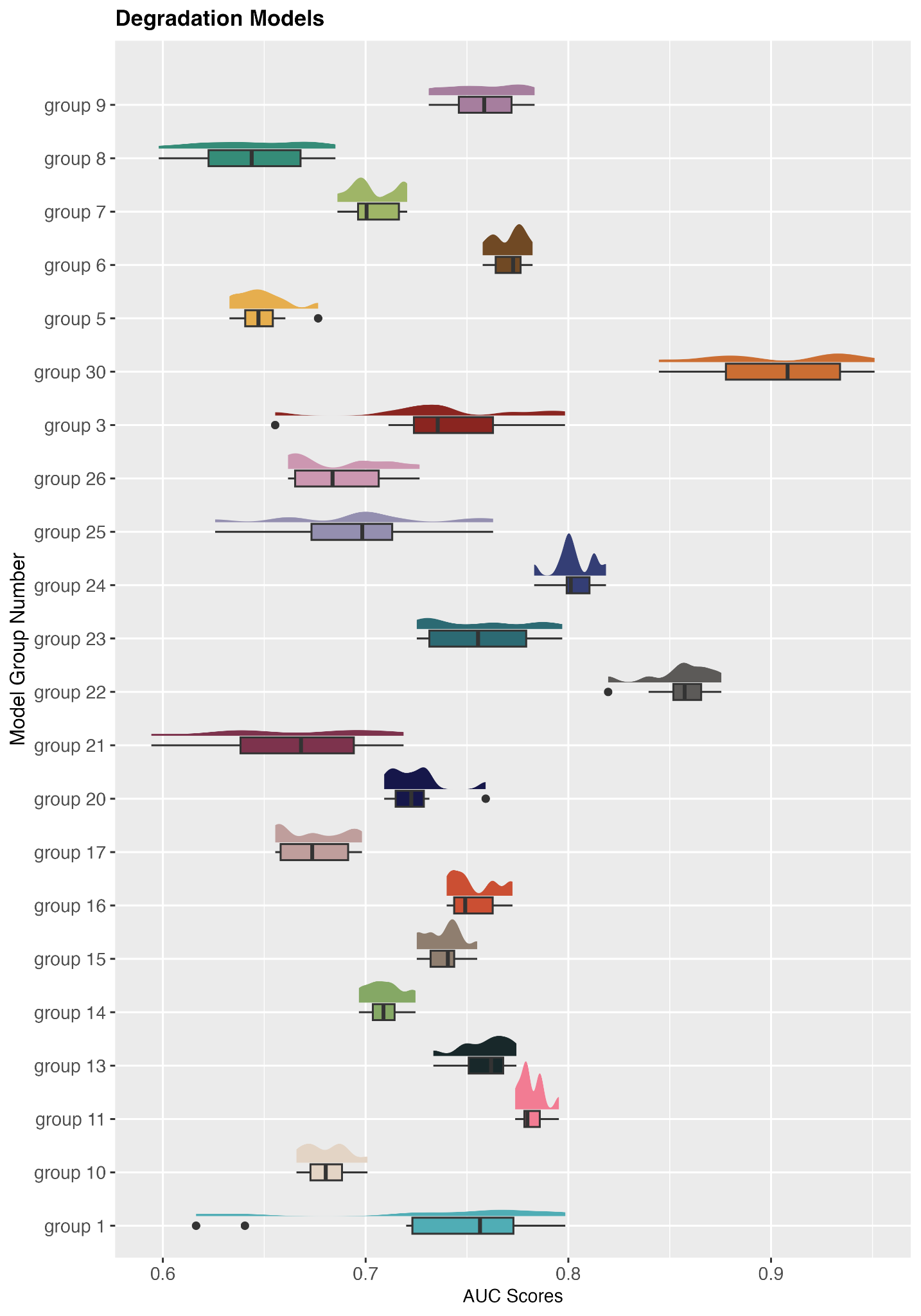

**Figure S2** | Distribution of area under the curve scores for (A) deforestation and (B) degradation random forest models. Ten individual models were built for each Key Biodiversity Area modelling group (n = 22) based on spatially disaggregated blocks. The predictions for each block were used to derive an ensemble mean probability, weighted by the AUC score of the block, for each model group.

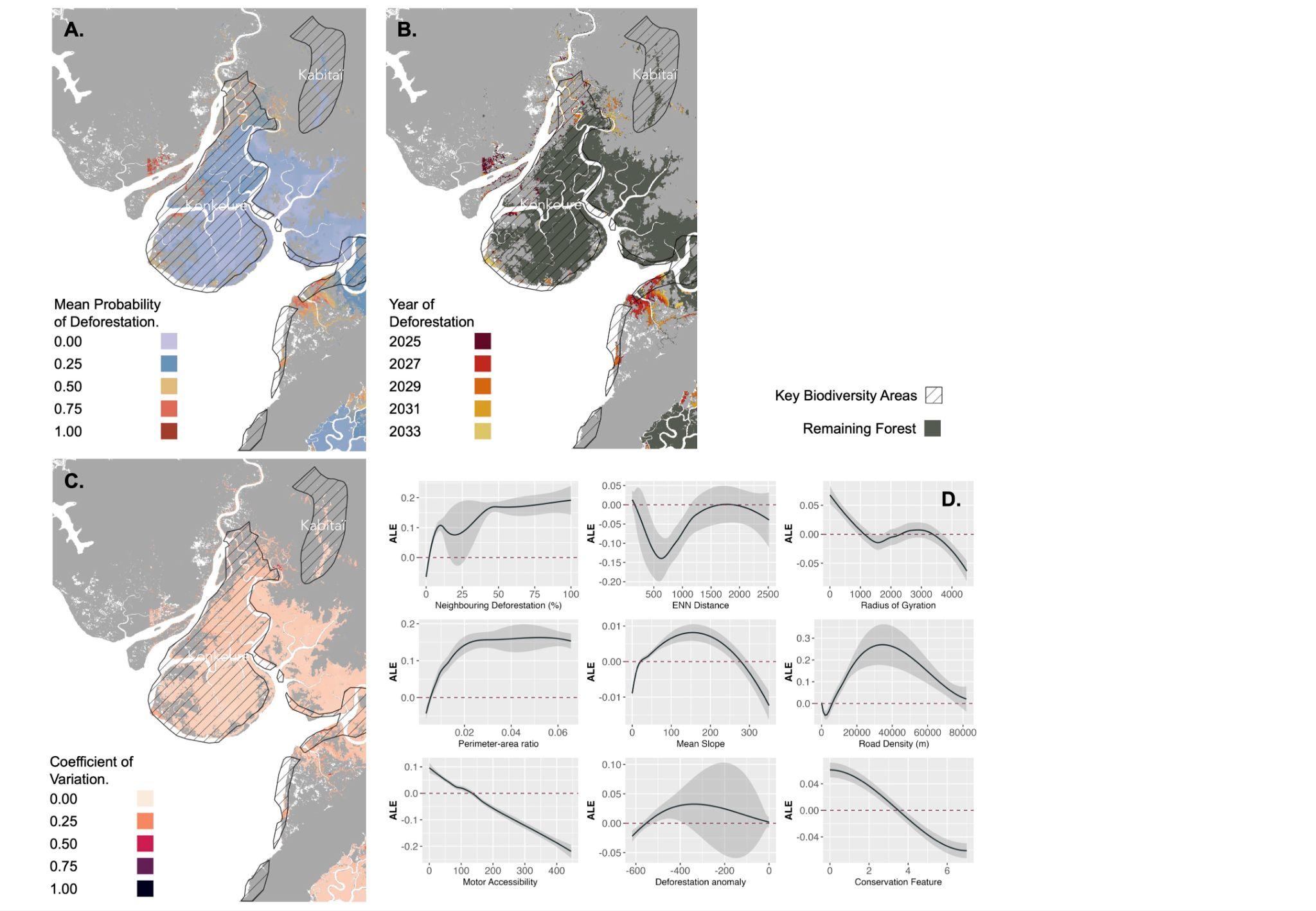

**Figure S3.1 |** Maps across the extent of model group 1, including Key Biodiversity Areas: Konkouré and Kabitaï, for **(A)** Mean probability of deforestation between 2025 and 2033 (inclusive), **(B)** Predicted deforestation event occurrence by year, **(C)** Mean coefficient of variation across model blocks (n = 10), and **(D)** Accumulated local effects plots for explanatory variables.

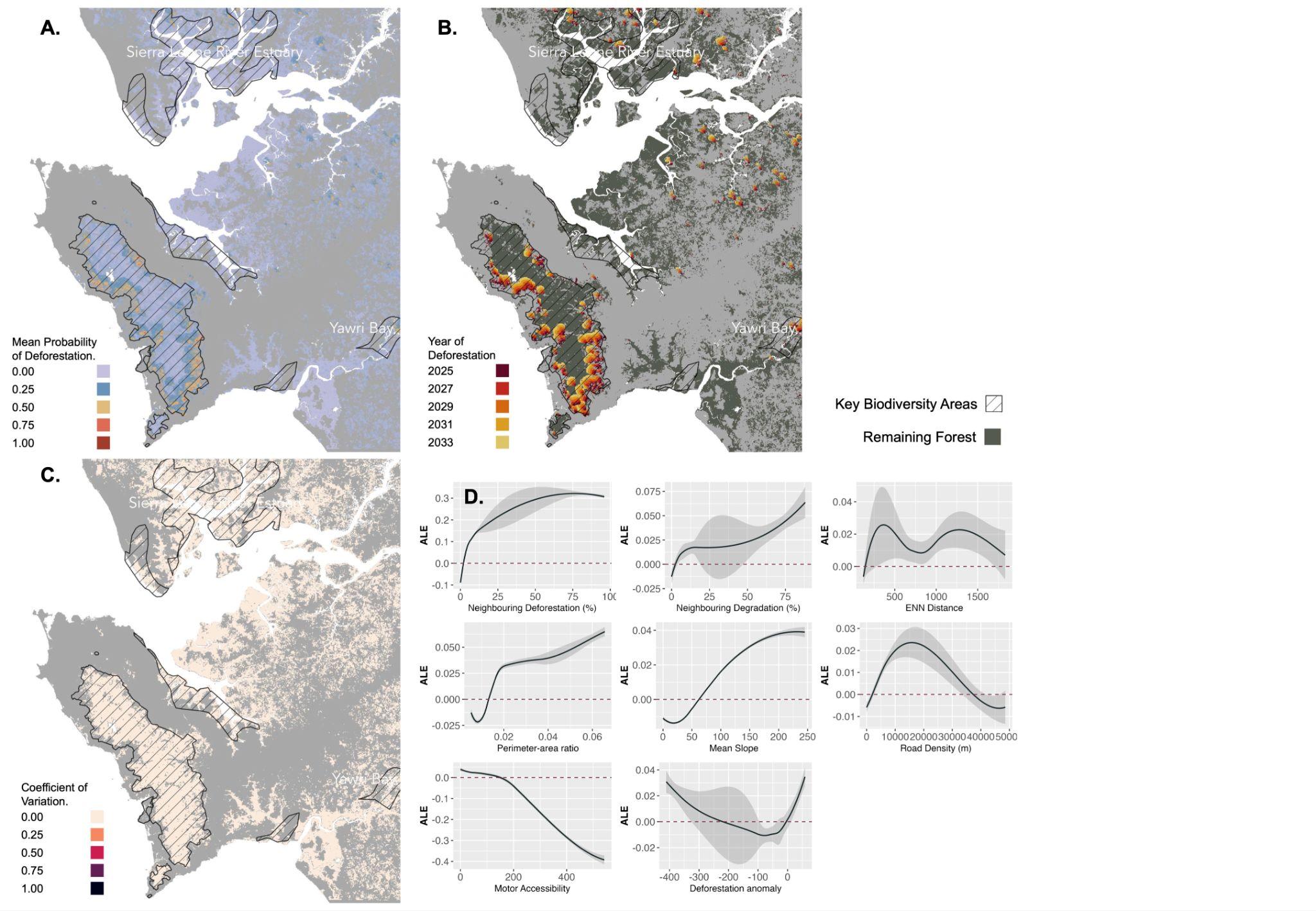

**Figure S3.2 |** Maps across the extent of model group 3, including Key Biodiversity Areas: Sierra Leone River Estuary, Western Area Peninsula Forest National Park and Yawri Bay, for **(A)** Mean probability of deforestation between 2025 and 2033 (inclusive), **(B)** Predicted deforestation event occurrence by year, **(C)** Mean coefficient of variation across model blocks (n = 10), and **(D)** Accumulated local effects plots for explanatory variables.

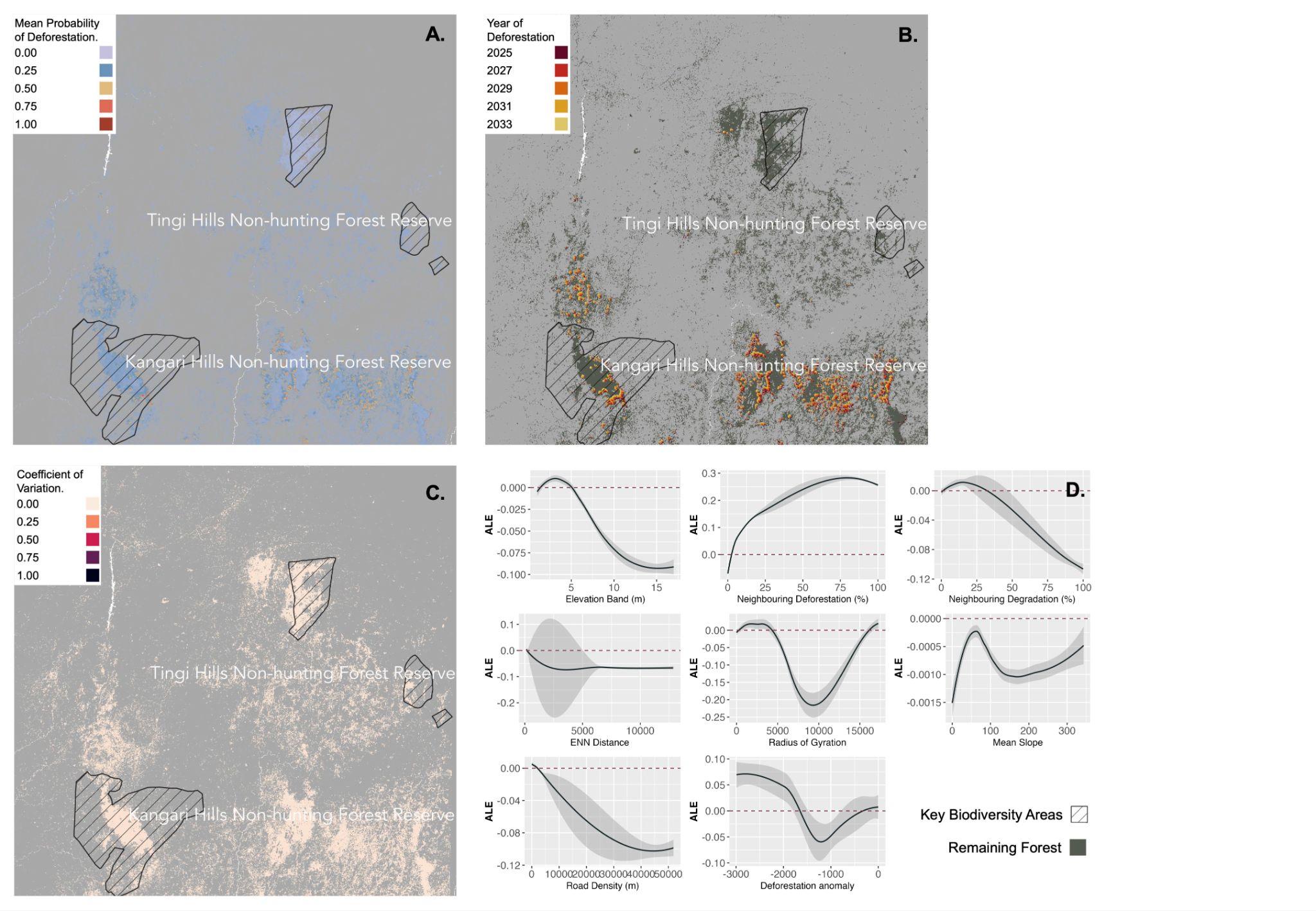

**Figure S3.3 |** Maps across the extent of model group 5, including Key Biodiversity Areas: Kangari Hills Non-hunting Forest Reserve, Loma Mountains Non-hunting Forest Reserve and Tingi Hills Non-hunting Forest Reserve, for **(A)** Mean probability of deforestation between 2025 and 2033 (inclusive), **(B)** Predicted deforestation event occurrence by year, **(C)** Mean coefficient of variation across model blocks (n = 10), and **(D)** Accumulated local effects plots for explanatory variables.

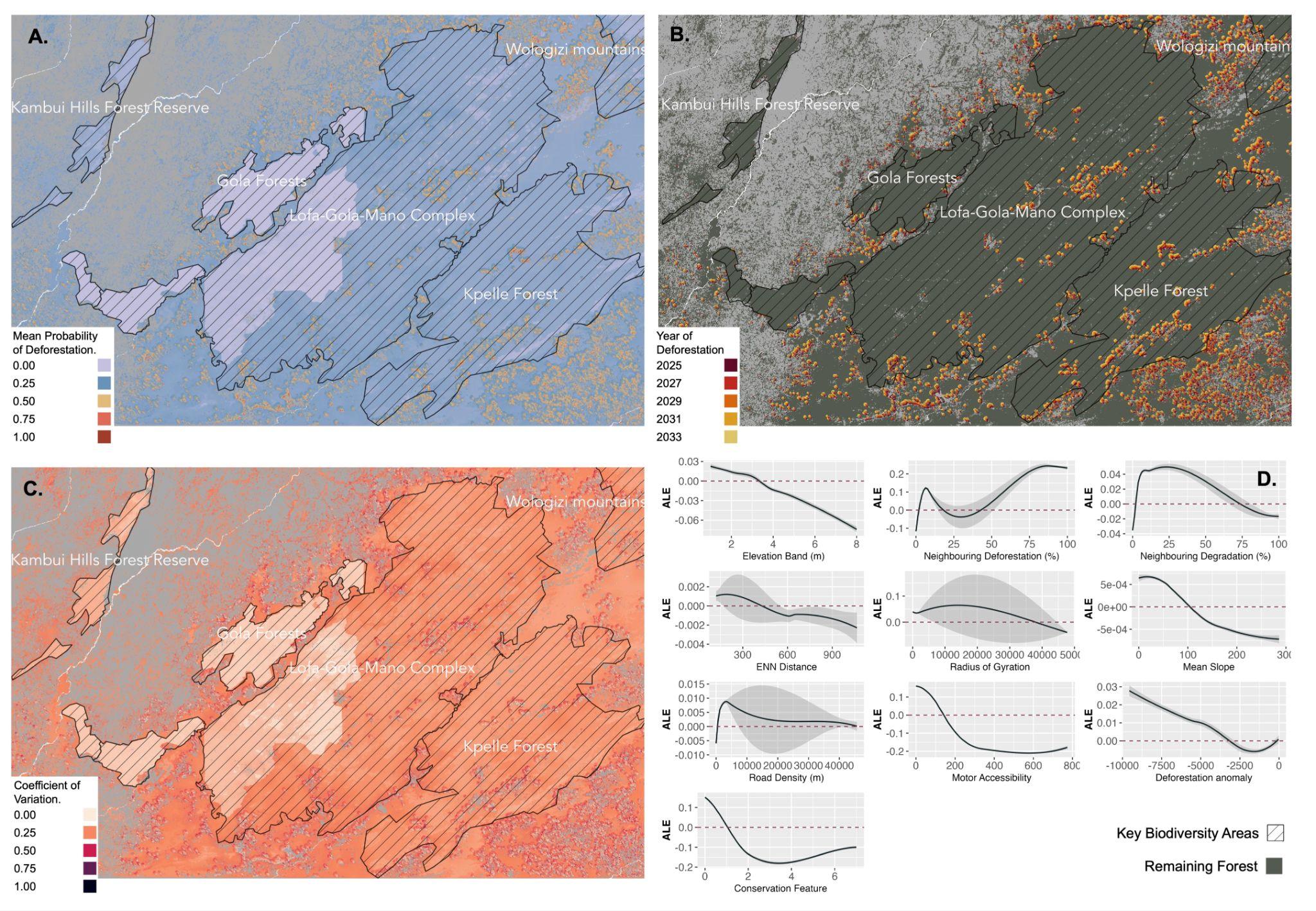

**Figure S3.4 |** Maps across the extent of model group 6, including Key Biodiversity Areas: Kambui Hill Forest Reserve, Gola Forests, Lofa-Gola-Mano Complex and Kpelle Forest, for **(A)** Mean probability of deforestation between 2025 and 2033 (inclusive), **(B)** Predicted deforestation event occurrence by year, **(C)** Mean coefficient of variation across model blocks (n = 10), and **(D)** Accumulated local effects plots for explanatory variables.

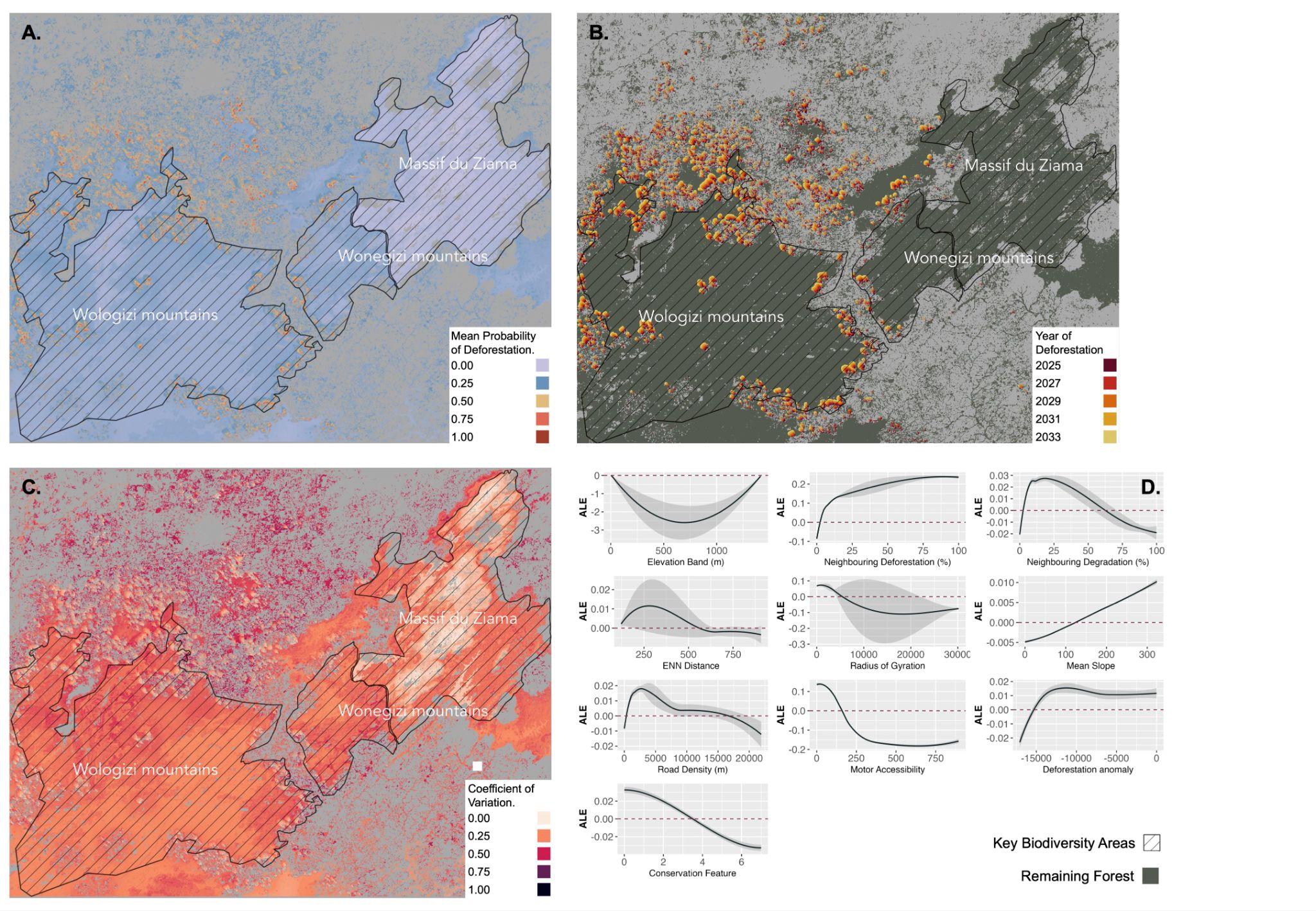

**Figure S3.5 |** Maps across the extent of model group 7, including Key Biodiversity Areas: Wologizi Mountains, Wonegizi Mountains and Massif du Ziama, for **(A)** Mean probability of deforestation between 2025 and 2033 (inclusive), **(B)** Predicted deforestation event occurrence by year, **(C)** Mean coefficient of variation across model blocks (n = 10), and **(D)** Accumulated local effects plots for explanatory variables.

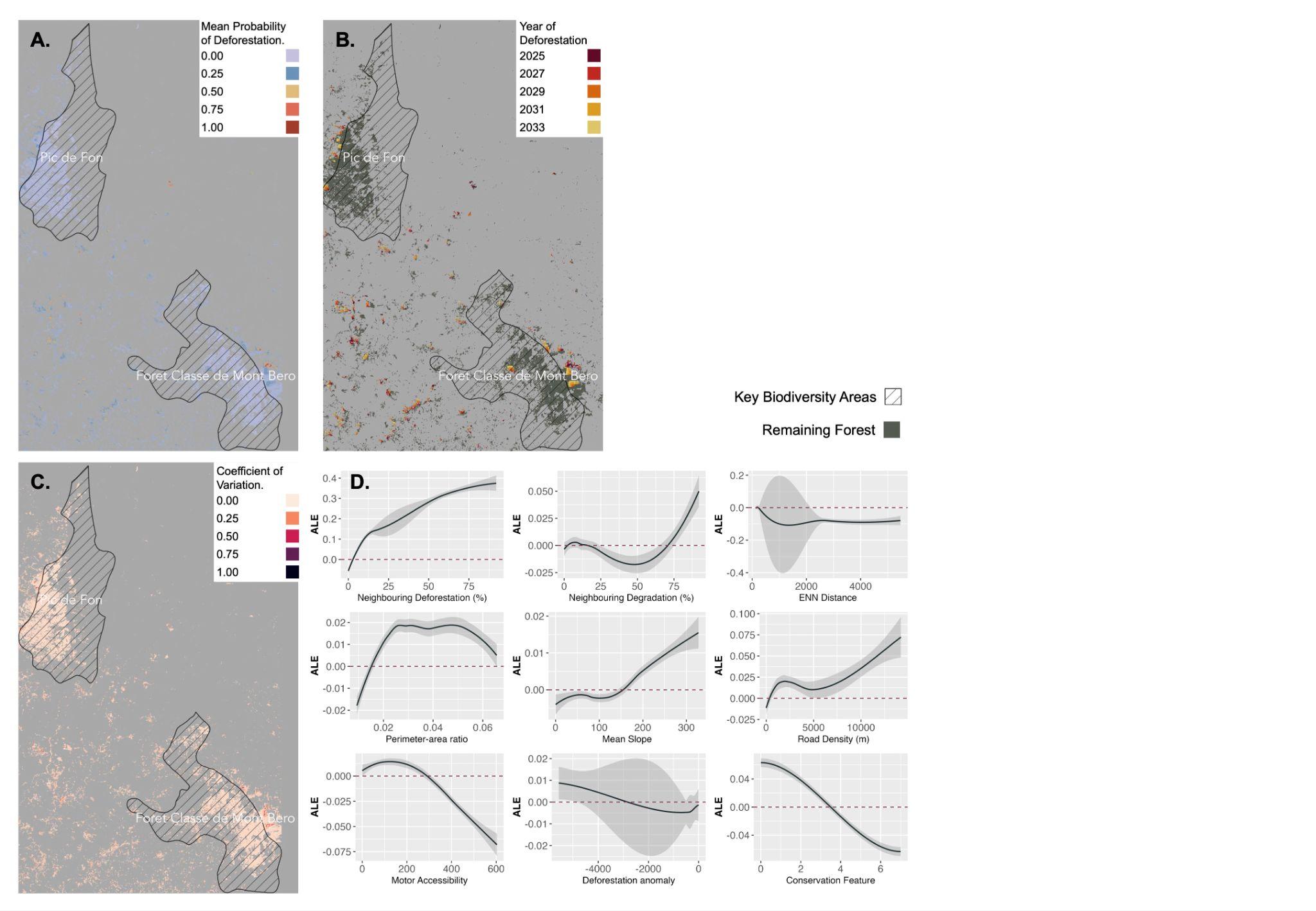

**Figure S3.6 |** Maps across the extent of model group 8, including Key Biodiversity Areas: Foret Classe de Mont Bero and Pic de Fon, for **(A)** Mean probability of deforestation between 2025 and 2033 (inclusive), **(B)** Predicted deforestation event occurrence by year, **(C)** Mean coefficient of variation across model blocks (n = 10), and **(D)** Accumulated local effects plots for explanatory variables.

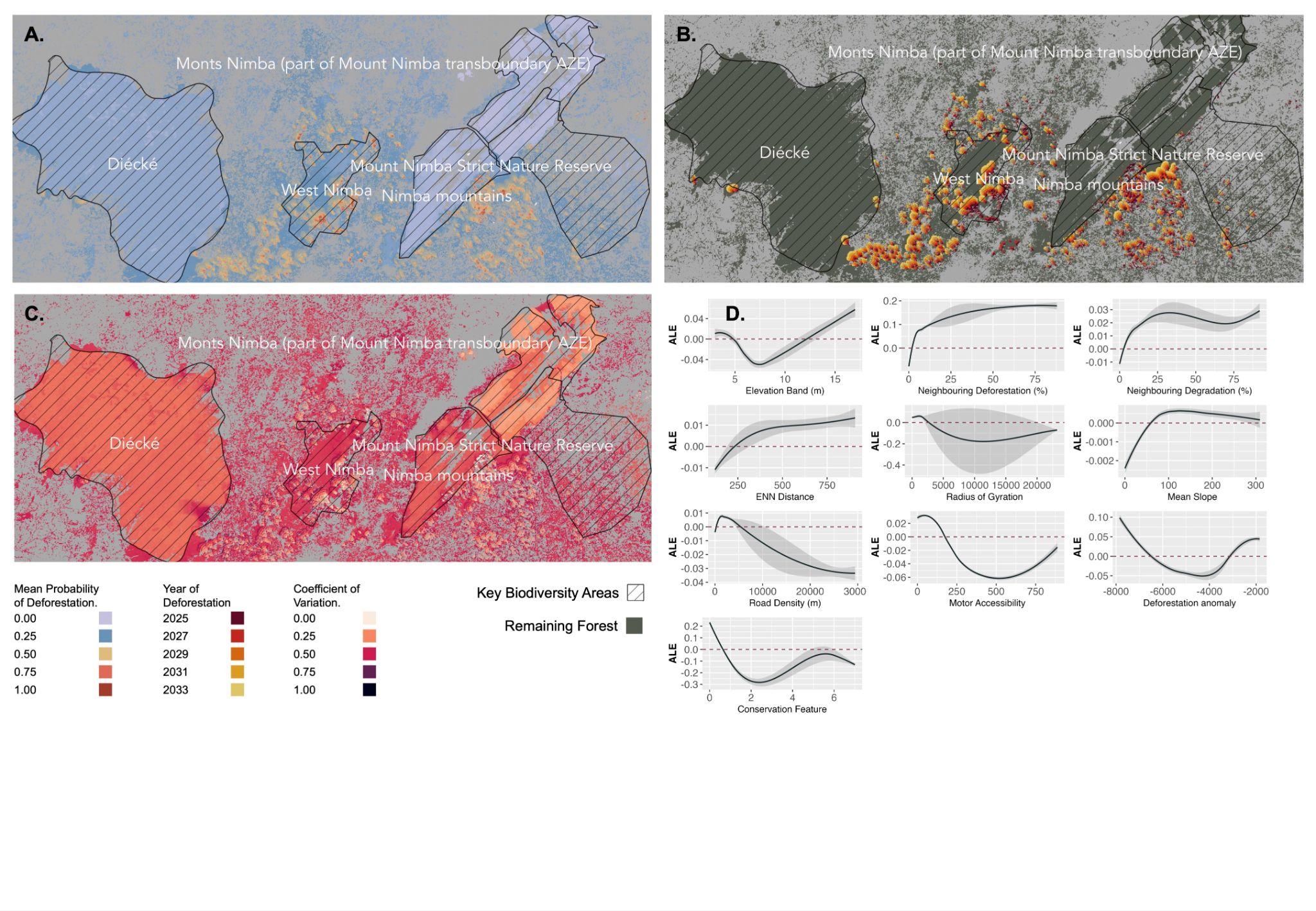

**Figure S3.7 |** Maps across the extent of model group 9, including Key Biodiversity Areas: Mount Nimba NR, West Nimba, Nimba Mountains, Monts Nimba and Diecke, for **(A)** Mean probability of deforestation between 2025 and 2033 (inclusive), **(B)** Predicted deforestation event occurrence by year, **(C)** Mean coefficient of variation across model blocks (n = 10), and **(D)** Accumulated local effects plots for explanatory variables.

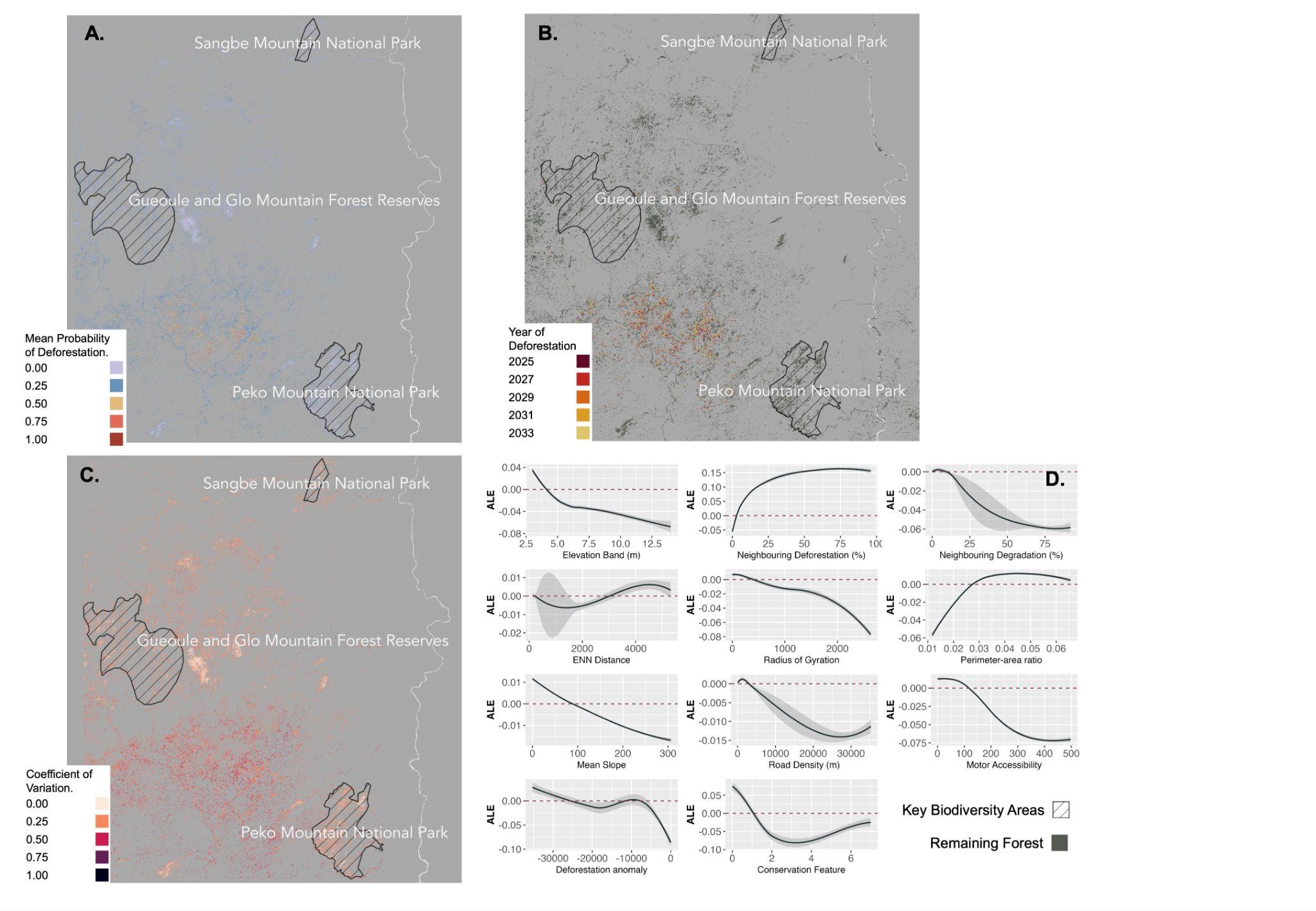

**Figure S3.8 |** Maps across the extent of model group 10, including Key Biodiversity Areas: Sangbe Mountain NP, Gueoule and Glo Mountain FR, and Peko Mountain NP for **(A)** Mean probability of deforestation between 2025 and 2033 (inclusive), **(B)** Predicted deforestation event occurrence by year, **(C)** Mean coefficient of variation across model blocks (n = 10), and **(D)** Accumulated local effects plots for explanatory variables.

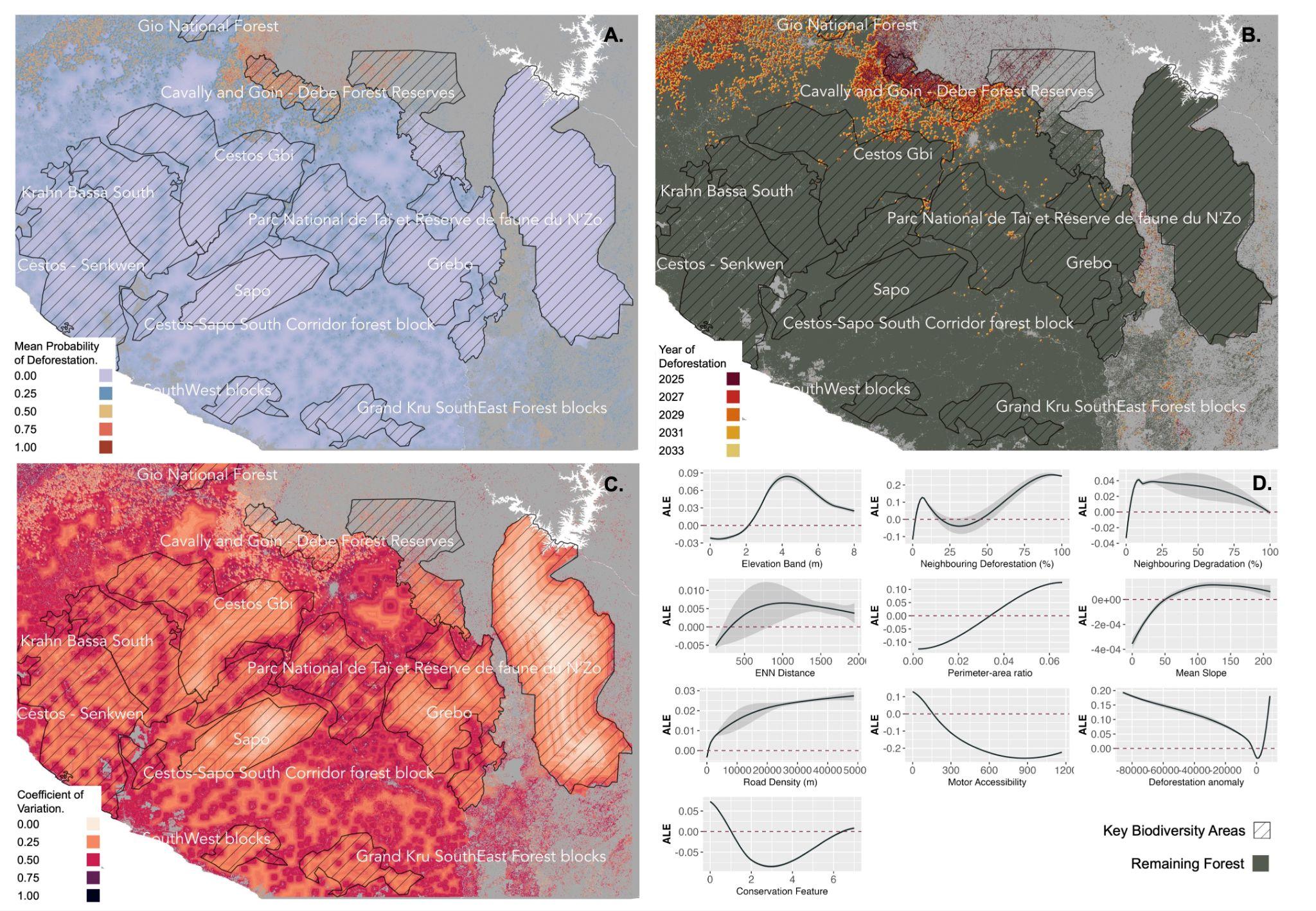

**Figure S3.9 |** Maps across the extent of model group 11, including Key Biodiversity Areas: Cavally and Going - Debe FRs, Zwedru, Gio NF, Sapo-Grebo Corridor, Grebo, Parc National de Tai et Reserve de faune du N’Zo, Cestos-Senkwen, Krahn Bassa South, Cestos-Sapo North Corridor, Gran Kru SouthEast, Grand Kru SouthWest, Sapo, and Weeni Creek for **(A)** Mean probability of deforestation between 2025 and 2033 (inclusive), **(B)** Predicted deforestation event occurrence by year, **(C)** Mean coefficient of variation across model blocks (n = 10), and **(D)** Accumulated local effects plots for explanatory variables.

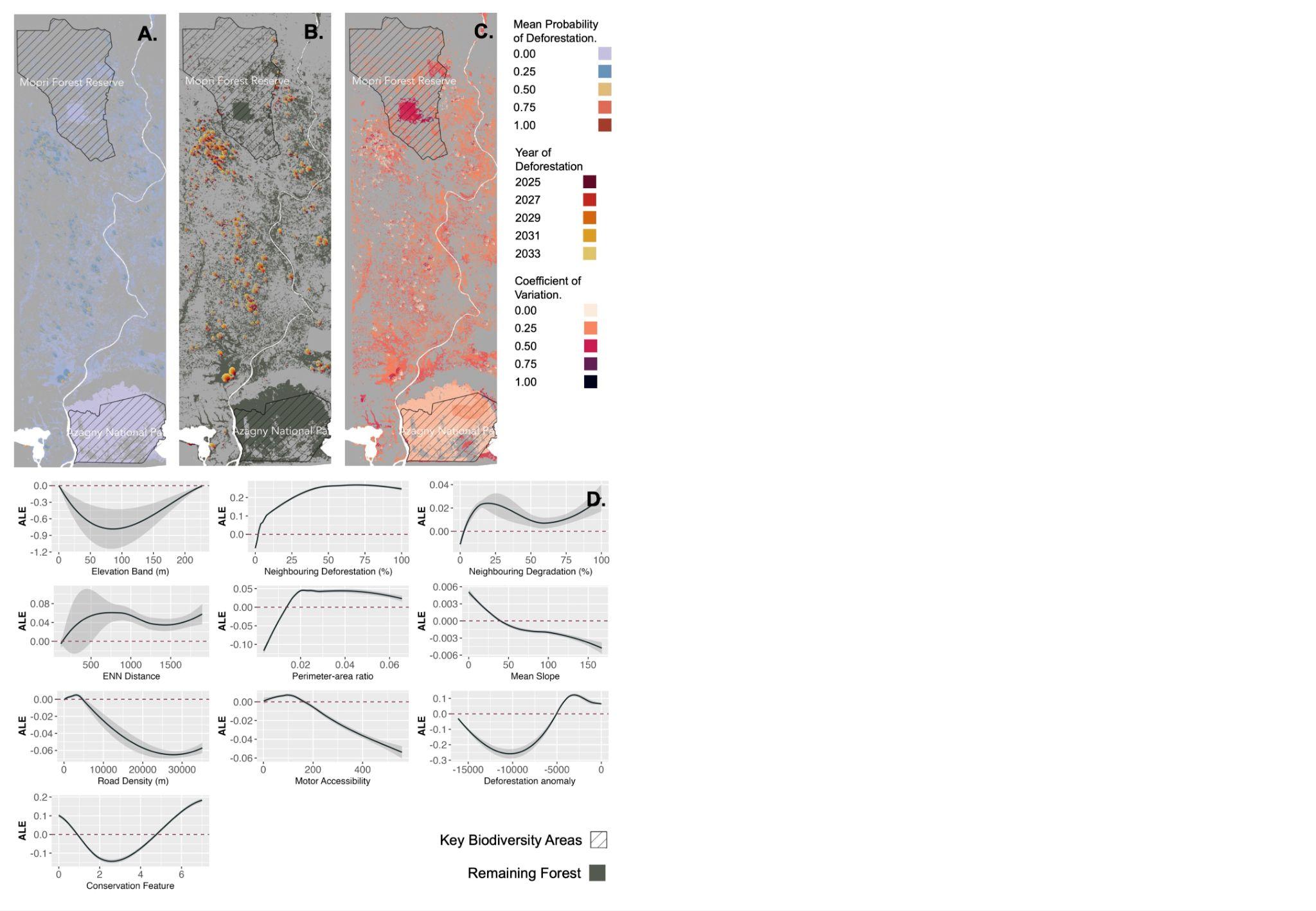

**Figure S3.10 |** Maps across the extent of model group 13, including Key Biodiversity Areas: Lamto Ecological Research Station, Mopri FR and Azagny NP, for **(A)** Mean probability of deforestation between 2025 and 2033 (inclusive), **(B)** Predicted deforestation event occurrence by year, **(C)** Mean coefficient of variation across model blocks (n = 10), and **(D)** Accumulated local effects plots for explanatory variables.

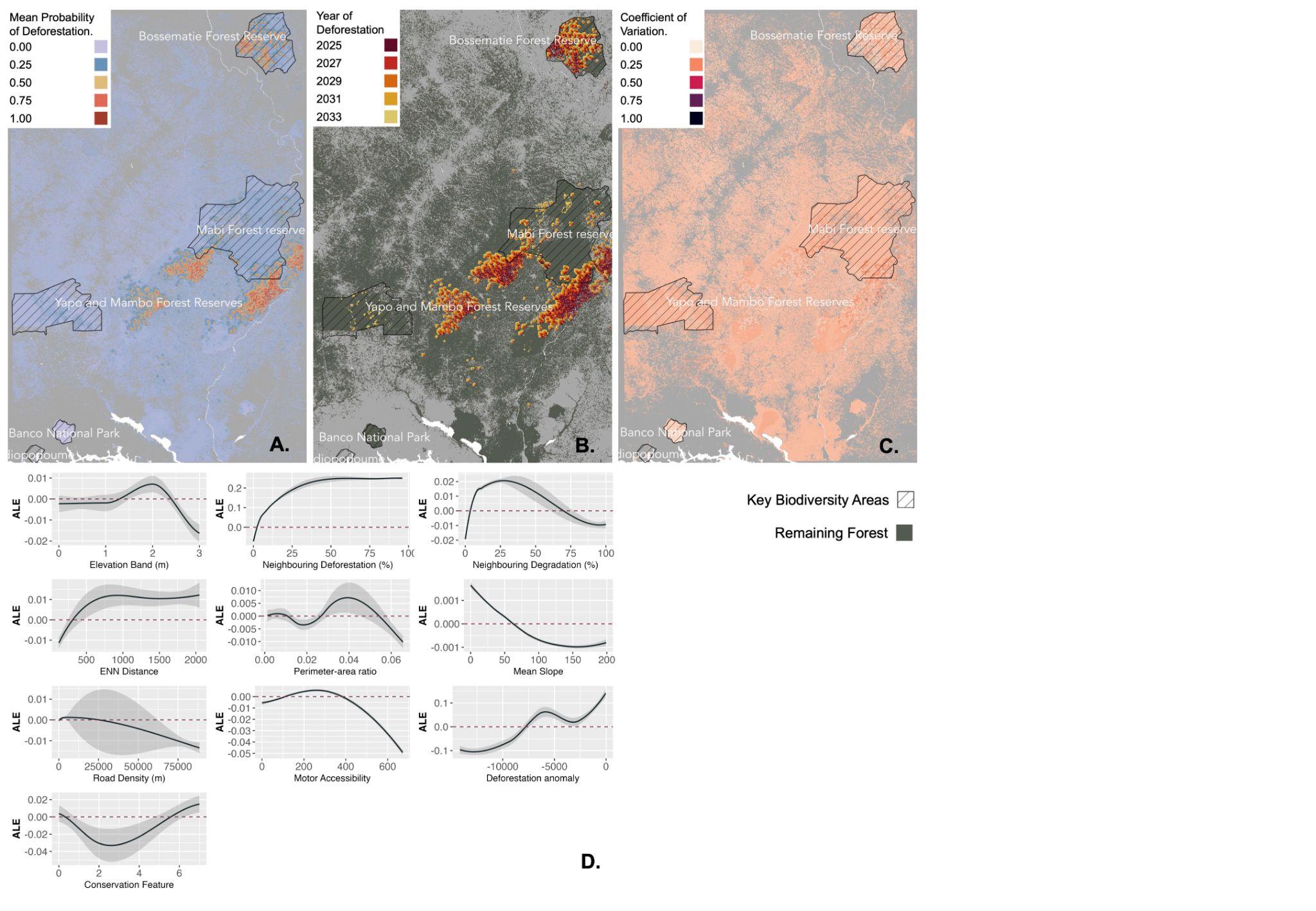

**Figure S3.11 |** Maps across the extent of model group 14, including Key Biodiversity Areas: Bossematie FR, Yapo and Mambo FR, Mabi FR, Adiopodoume and Banco NP, for **(A)** Mean probability of deforestation between 2025 and 2033 (inclusive), **(B)** Predicted deforestation event occurrence by year, **(C)** Mean coefficient of variation across model blocks (n = 10), and **(D)** Accumulated local effects plots for explanatory variables.

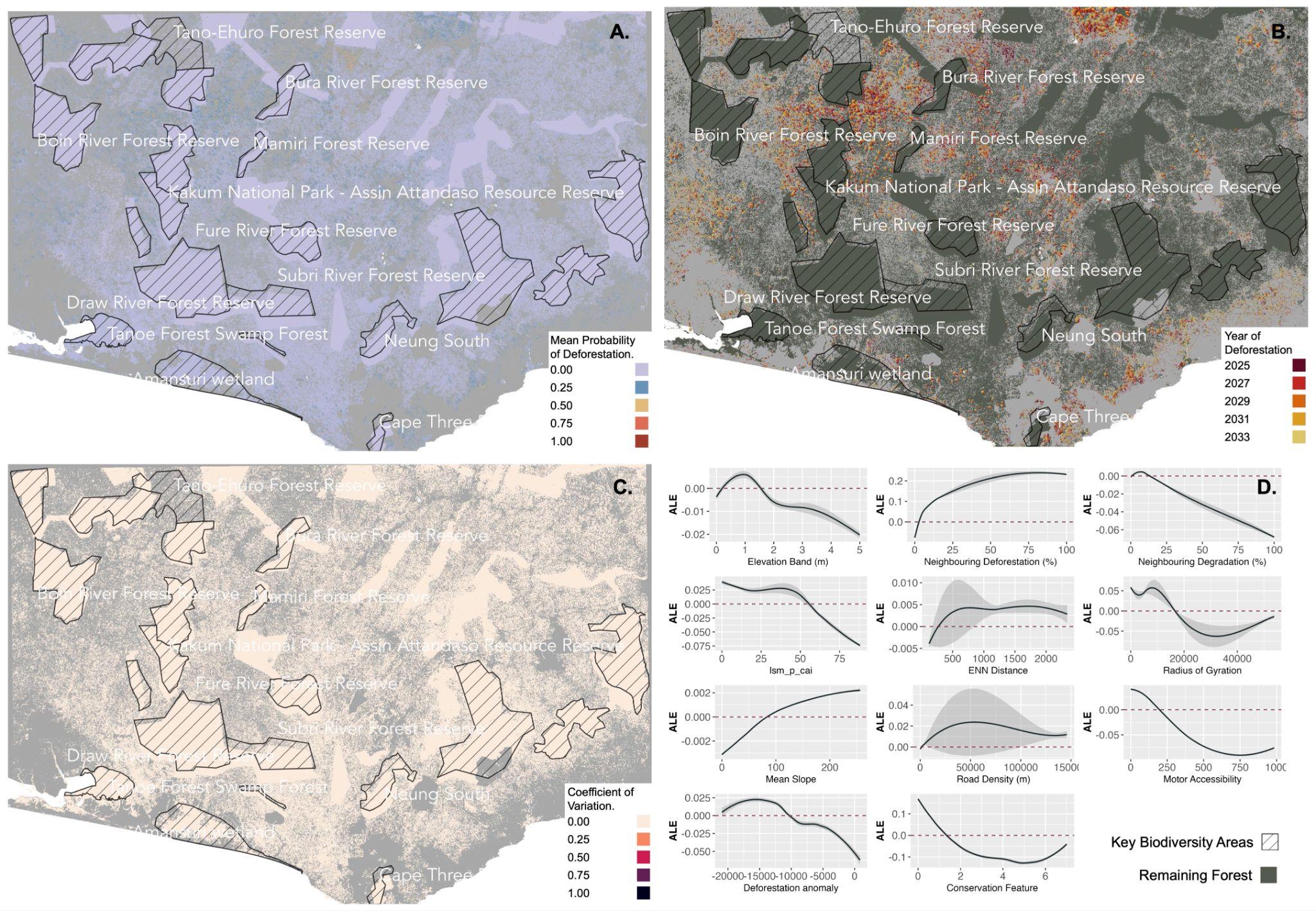

**Figure S3.12 |** Maps across the extent of model group 15, including Key Biodiversity Areas: Amansuri wetland, Ebi River Shelterbelt FR, Tanoe Forest, Jema-Asemkrom FR, Neung South, Tano-Nimiri FR, Ankasa Resource Reserve, Mamiri FR, Kakum NP, Subri River FR, Yoyo River FR, Pra-Sushien FR, Tano-Anwia FR, Draw River FR, Tano-Ehuro FR, Cape Three Points FR, Boin Tano FR, Bura River FR, Boin River FR, Dadieso FR, and Fure River FR for **(A)** Mean probability of deforestation between 2025 and 2033 (inclusive), **(B)** Predicted deforestation event occurrence by year, **(C)** Mean coefficient of variation across model blocks (n = 10), and **(D)** Accumulated local effects plots for explanatory variables.

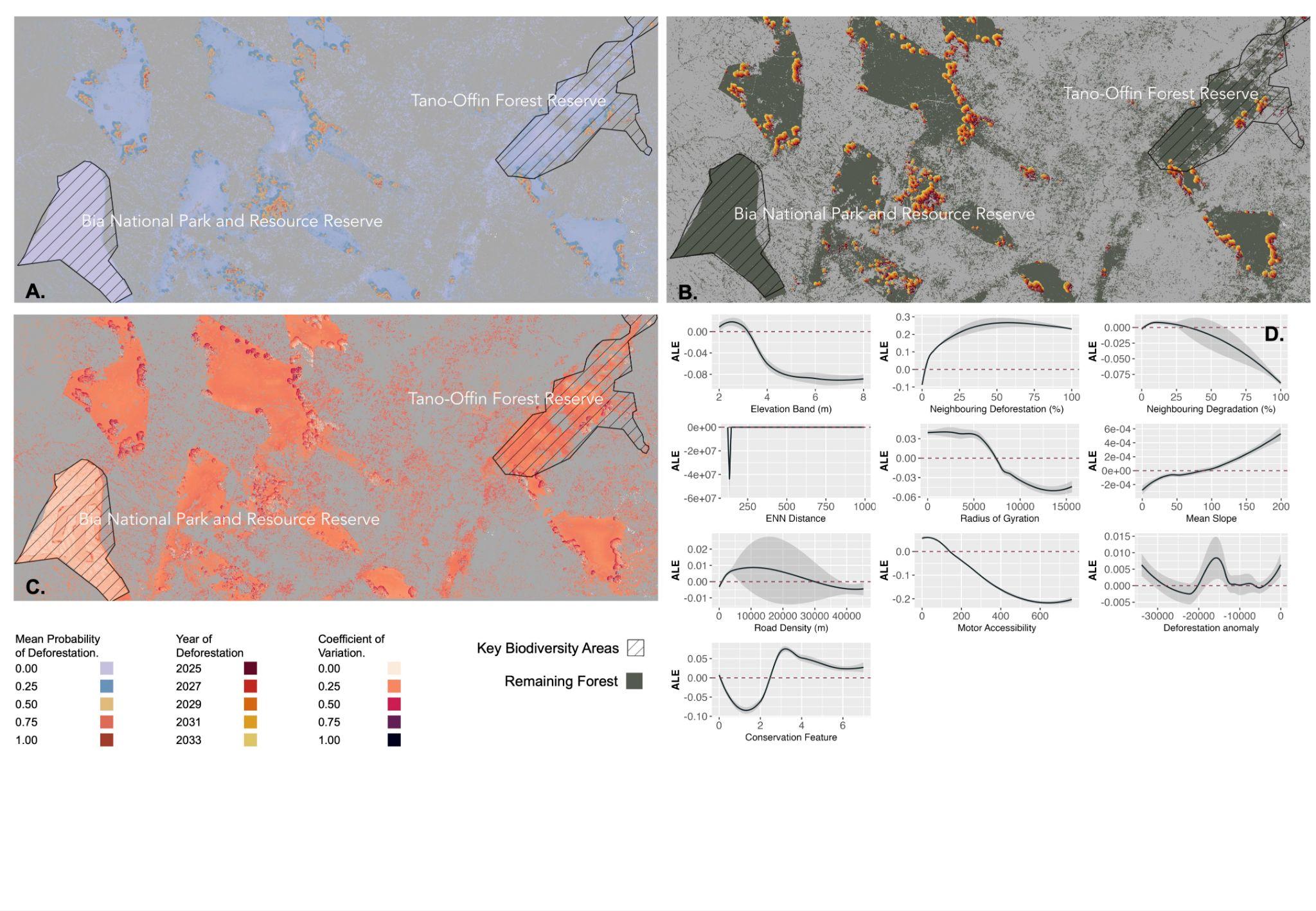

**Figure S3.13 |** Maps across the extent of model group 16, including Key Biodiversity Areas: Tano-Offin FR and Bia NP for **(A)** Mean probability of deforestation between 2025 and 2033 (inclusive), **(B)** Predicted deforestation event occurrence by year, **(C)** Mean coefficient of variation across model blocks (n = 10), and **(D)** Accumulated local effects plots for explanatory variables.

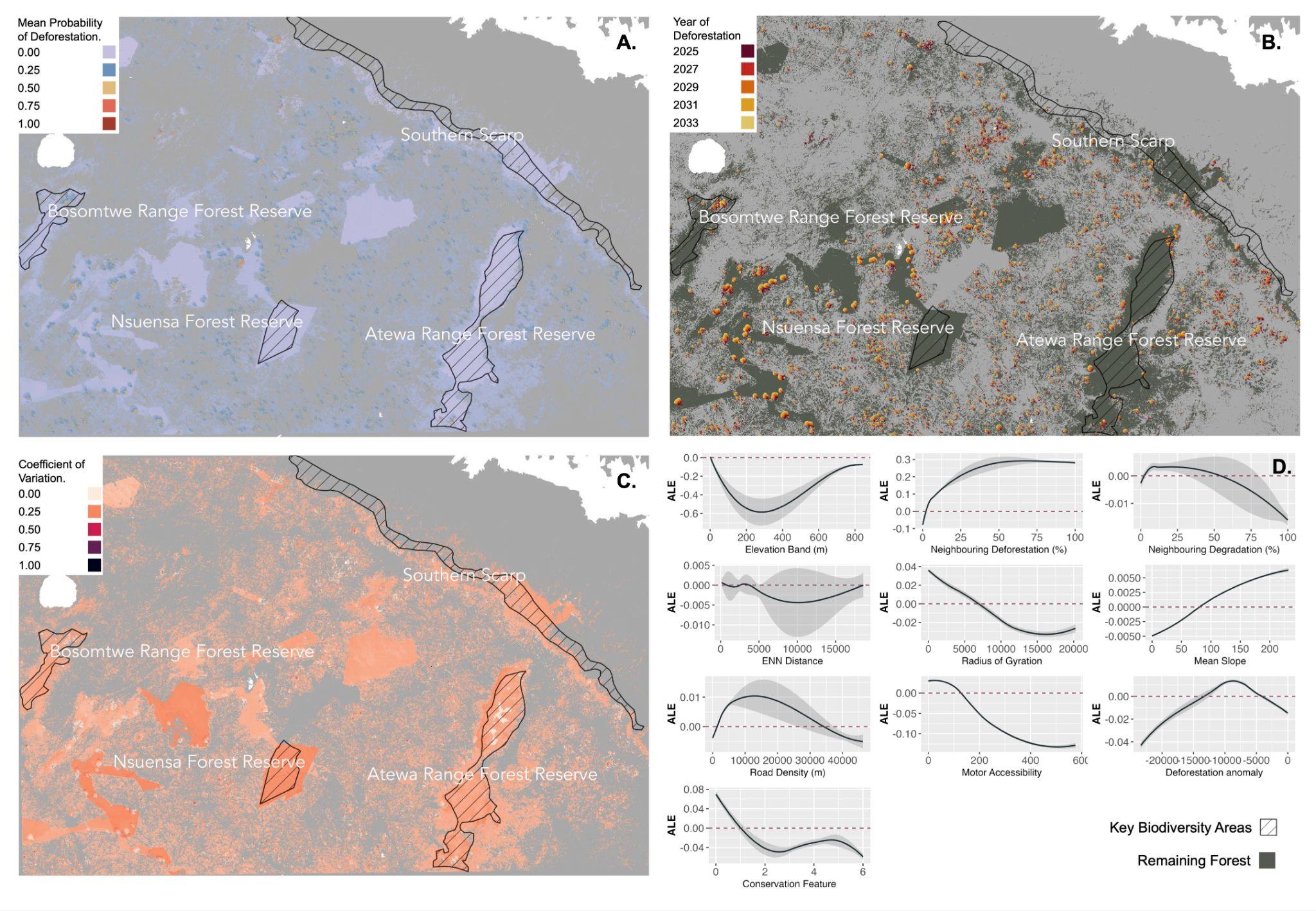

**Figure S3.14 |** Maps across the extent of model group 17, including Key Biodiversity Areas: Atewa Range FR, Bosomtwe Range FR, Southern Scarp, Sapawsu FR, and Nsuensa FR for **(A)** Mean probability of deforestation between 2025 and 2033 (inclusive), **(B)** Predicted deforestation event occurrence by year, **(C)** Mean coefficient of variation across model blocks (n = 10), and **(D)** Accumulated local effects plots for explanatory variables.

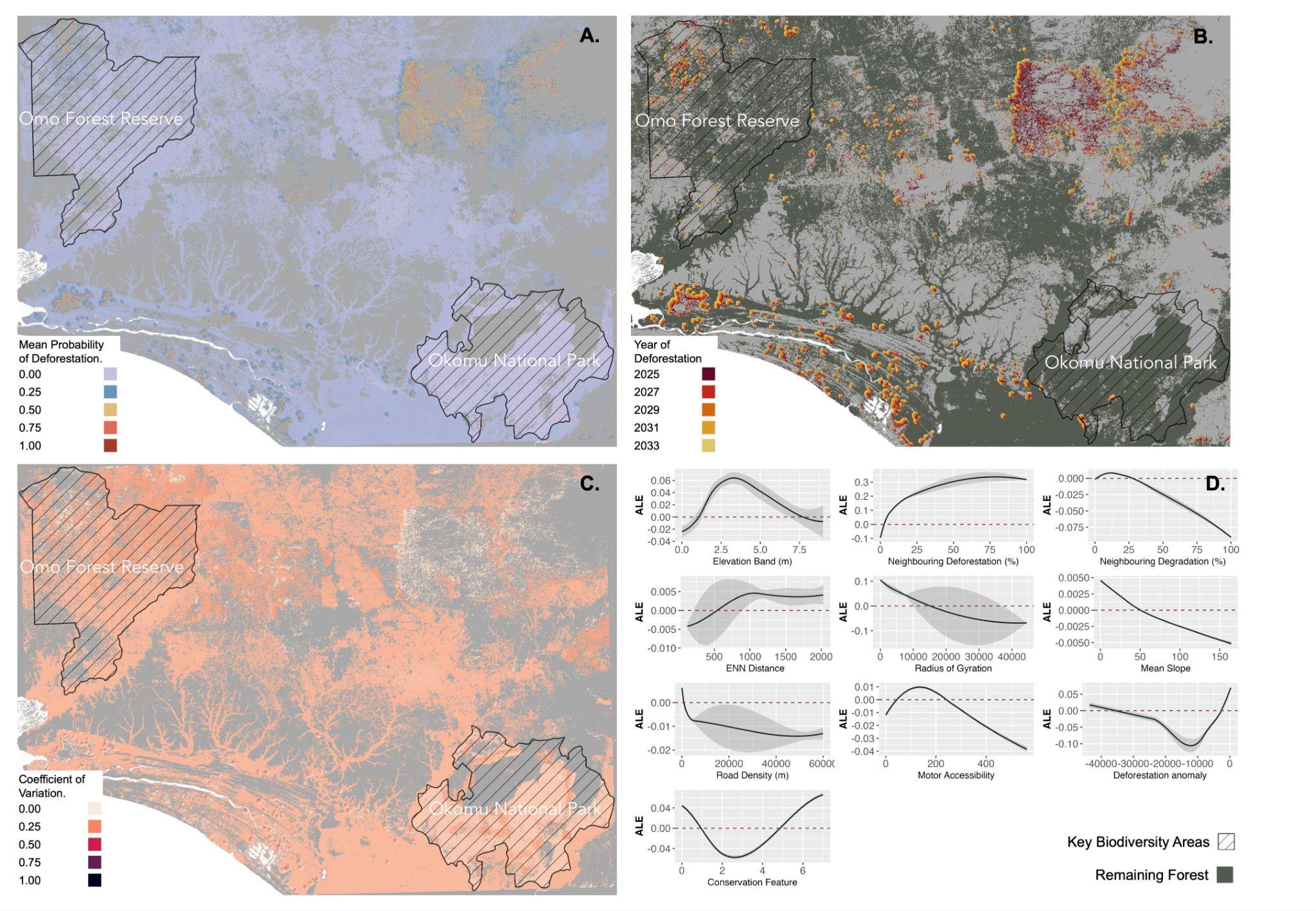

**Figure S3.15 |** Maps across the extent of model group 20, including Key Biodiversity Areas: Omo FR and Okomu NP, for **(A)** Mean probability of deforestation between 2025 and 2033 (inclusive), **(B)** Predicted deforestation event occurrence by year, **(C)** Mean coefficient of variation across model blocks (n = 10), and **(D)** Accumulated local effects plots for explanatory variables.

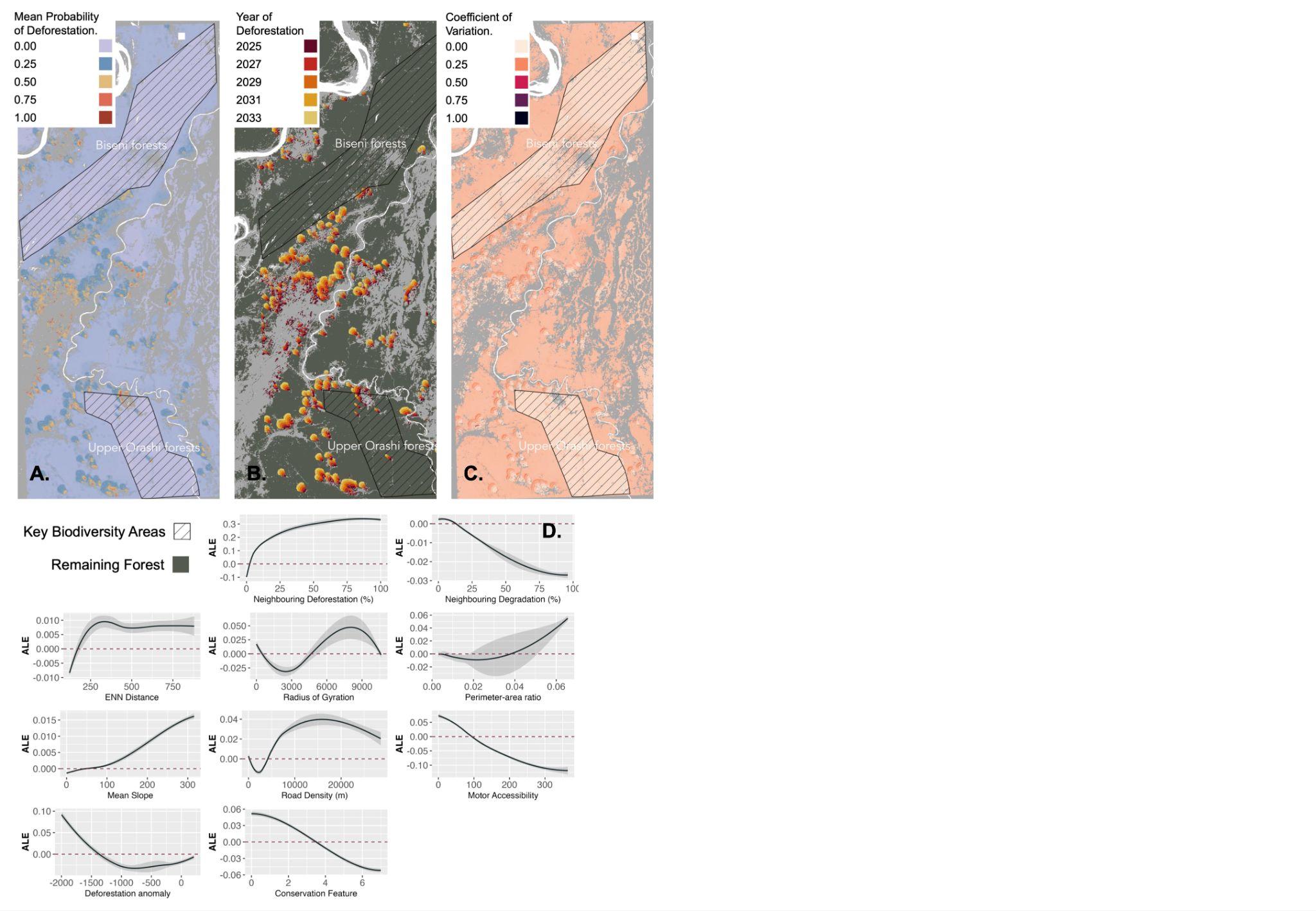

**Figure S3.16 |** Maps across the extent of model group 21, including Key Biodiversity Areas: XXXX, for **(A)** Mean probability of deforestation between 2025 and 2033 (inclusive), **(B)** Predicted deforestation event occurrence by year, **(C)** Mean coefficient of variation across model blocks (n = 10), and **(D)** Accumulated local effects plots for explanatory variables

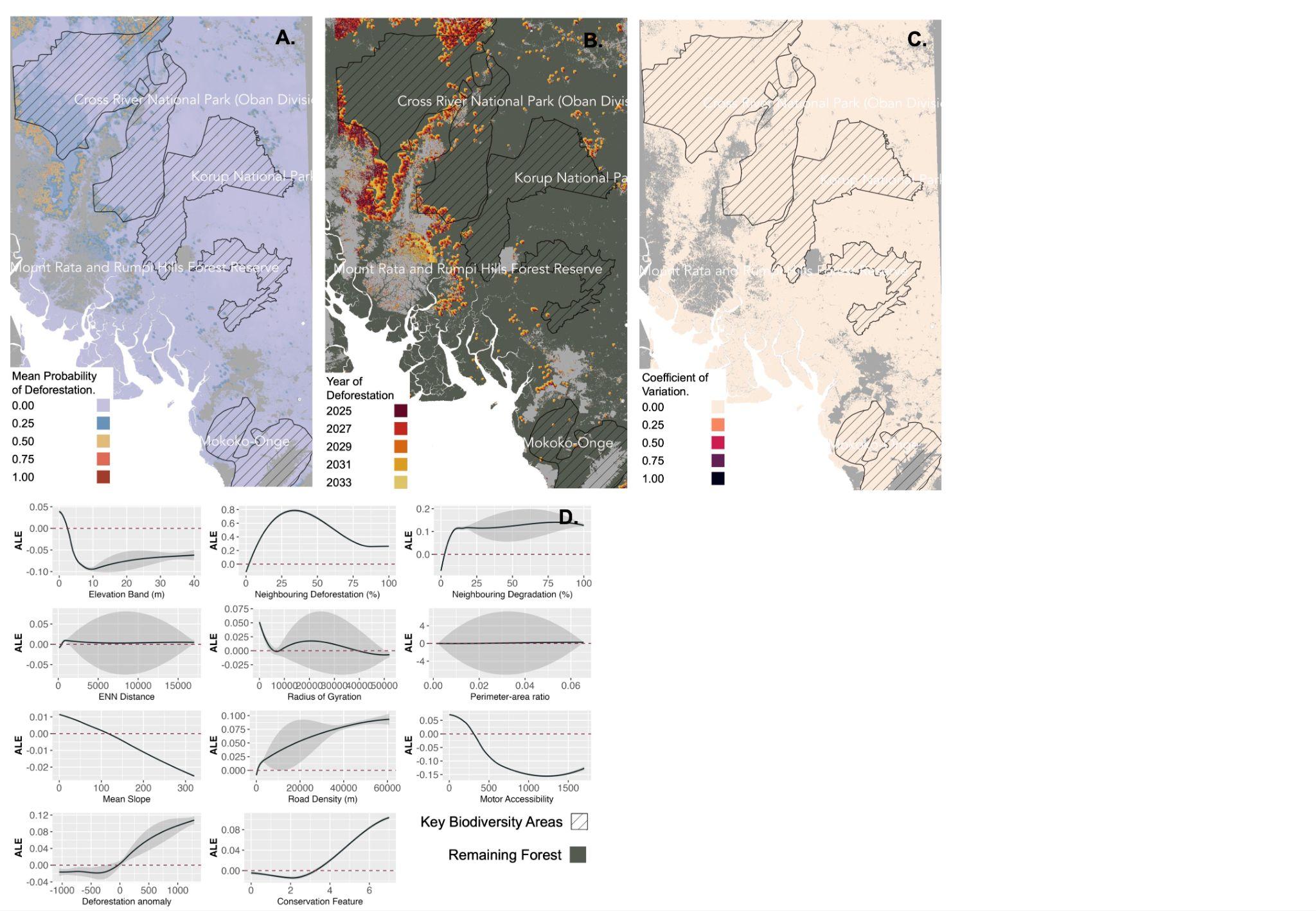

**Figure S3.17 |** Maps across the extent of model group 22, including Key Biodiversity Areas: Cross River NP, Mount Cameroon and Mokoko-Onge, Korup NP and Mount Rata and Rumpi Hills FR for **(A)** Mean probability of deforestation between 2025 and 2033 (inclusive), **(B)** Predicted deforestation event occurrence by year, **(C)** Mean coefficient of variation across model blocks (n = 10), and **(D)** Accumulated local effects plots for explanatory variables.

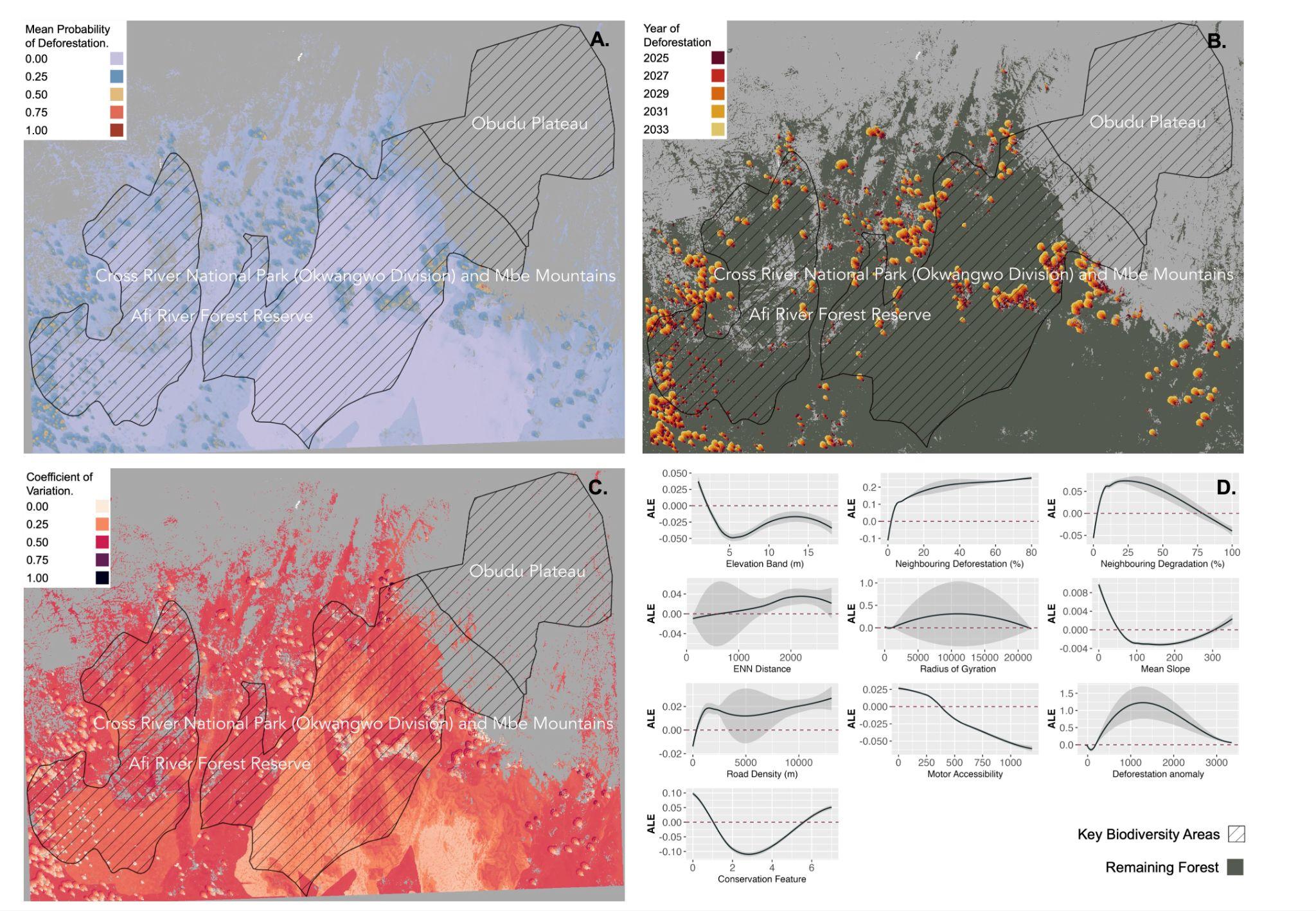

**Figure S3.18 |** Maps across the extent of model group 23, including Key Biodiversity Areas: Obudu Plateau, Afi River FR and Cross River NP and Mbe Mountains for **(A)** Mean probability of deforestation between 2025 and 2033 (inclusive), **(B)** Predicted deforestation event occurrence by year, **(C)** Mean coefficient of variation across model blocks (n = 10), and **(D)** Accumulated local effects plots for explanatory variables.

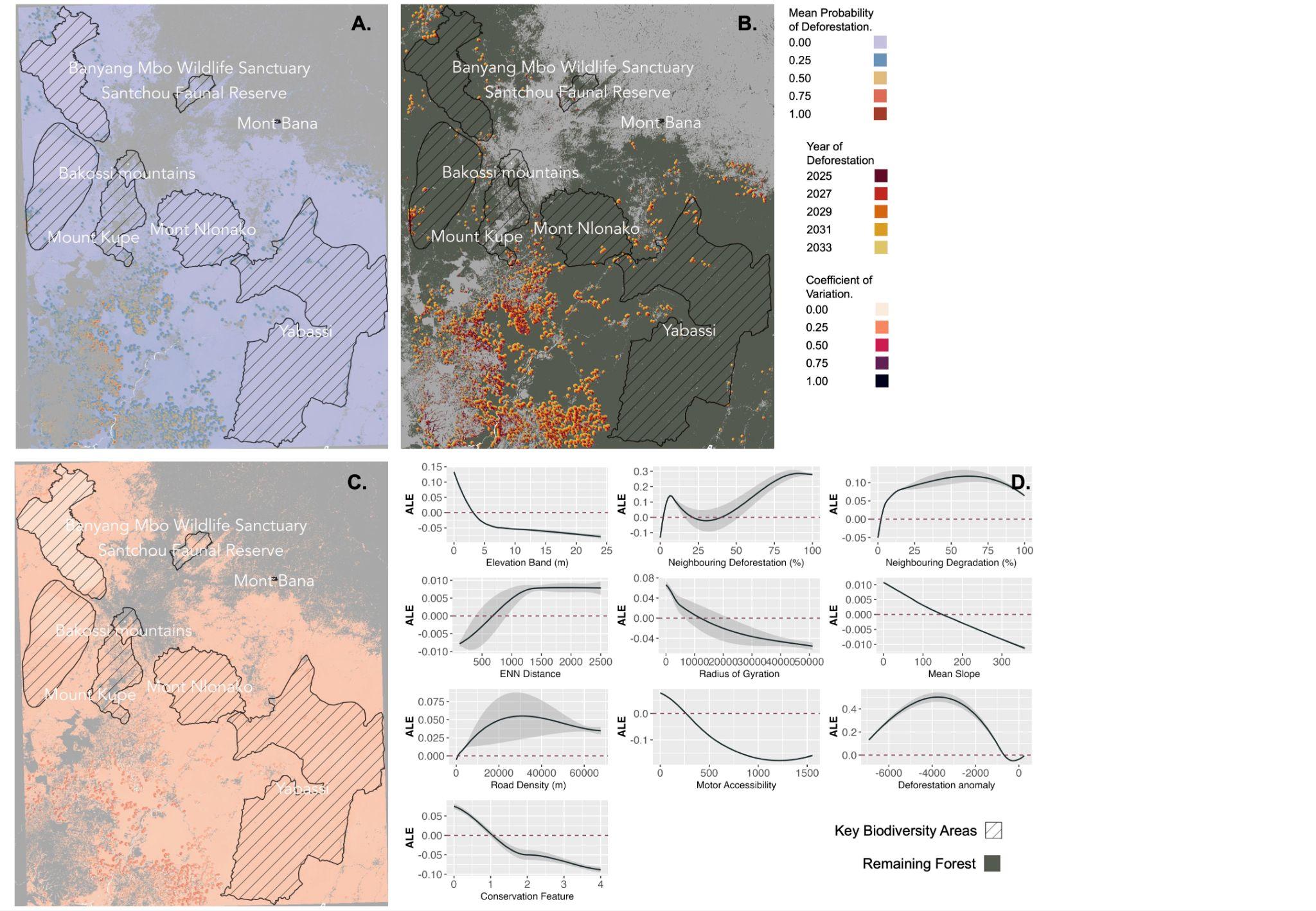

**Figure S3.19 |** Maps across the extent of model group 24, including Key Biodiversity Areas: Santchou Faunal Reserve, Eastern Bamenda highlands, Mont Manengouba, Mont Nlonako, Bakossi mountains, Yabassi, Banyang Mbo Wildlife Sanctuary, Mont Bana, and Mount Kupe for **(A)** Mean probability of deforestation between 2025 and 2033 (inclusive), **(B)** Predicted deforestation event occurrence by year, **(C)** Mean coefficient of variation across model blocks (n = 10), and **(D)** Accumulated local effects plots for explanatory variables.

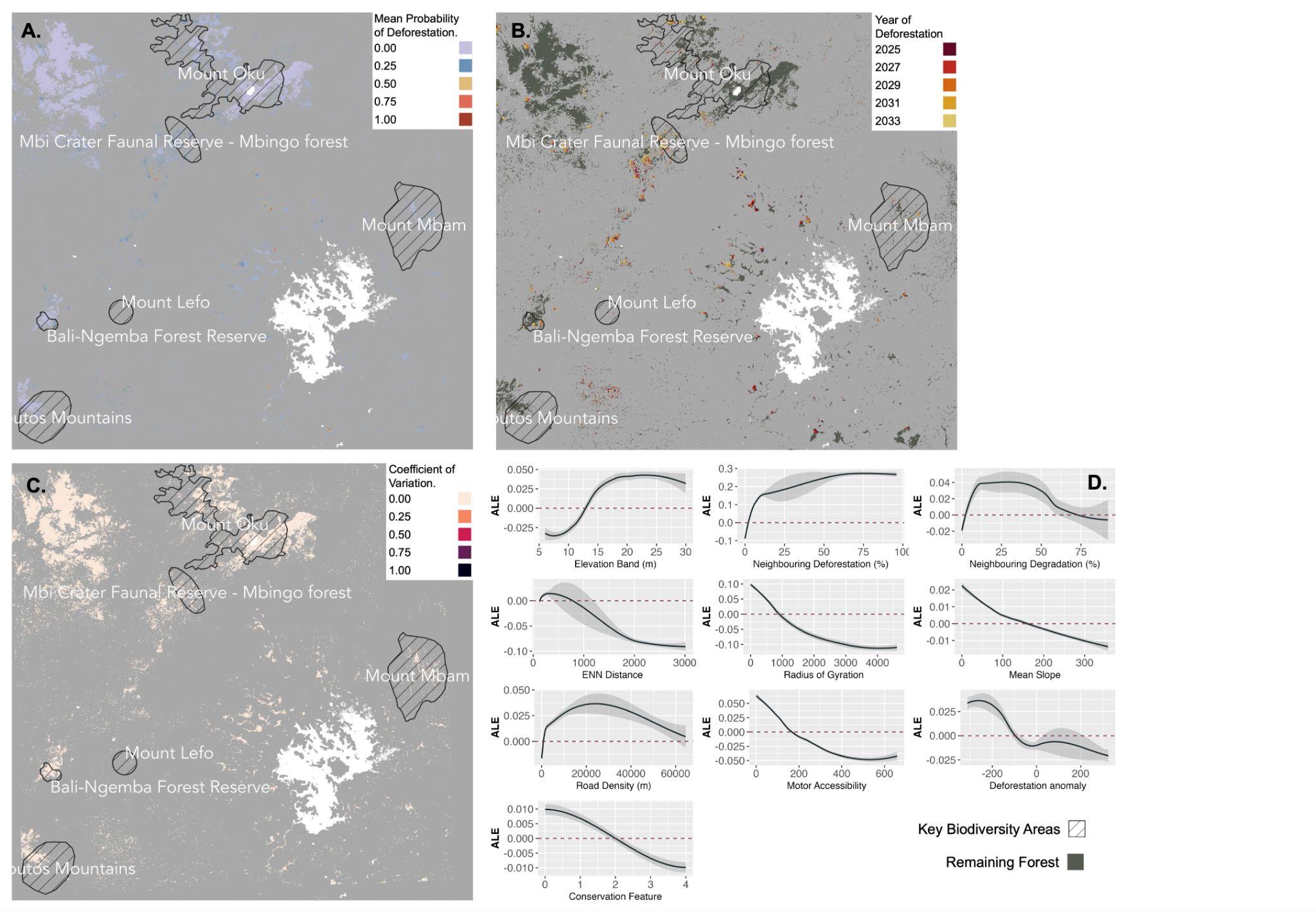

**Figure S3.20 |** Maps across the extent of model group 25, including Key Biodiversity Areas: Mount Oku, Mbi Crater Faunal Reserve, Bali-Ngemba FR, Bamboutos Mountains, Mount Mbam, and Mount Lefo for **(A)** Mean probability of deforestation between 2025 and 2033 (inclusive), **(B)** Predicted deforestation event occurrence by year, **(C)** Mean coefficient of variation across model blocks (n = 10), and **(D)** Accumulated local effects plots for explanatory variables.

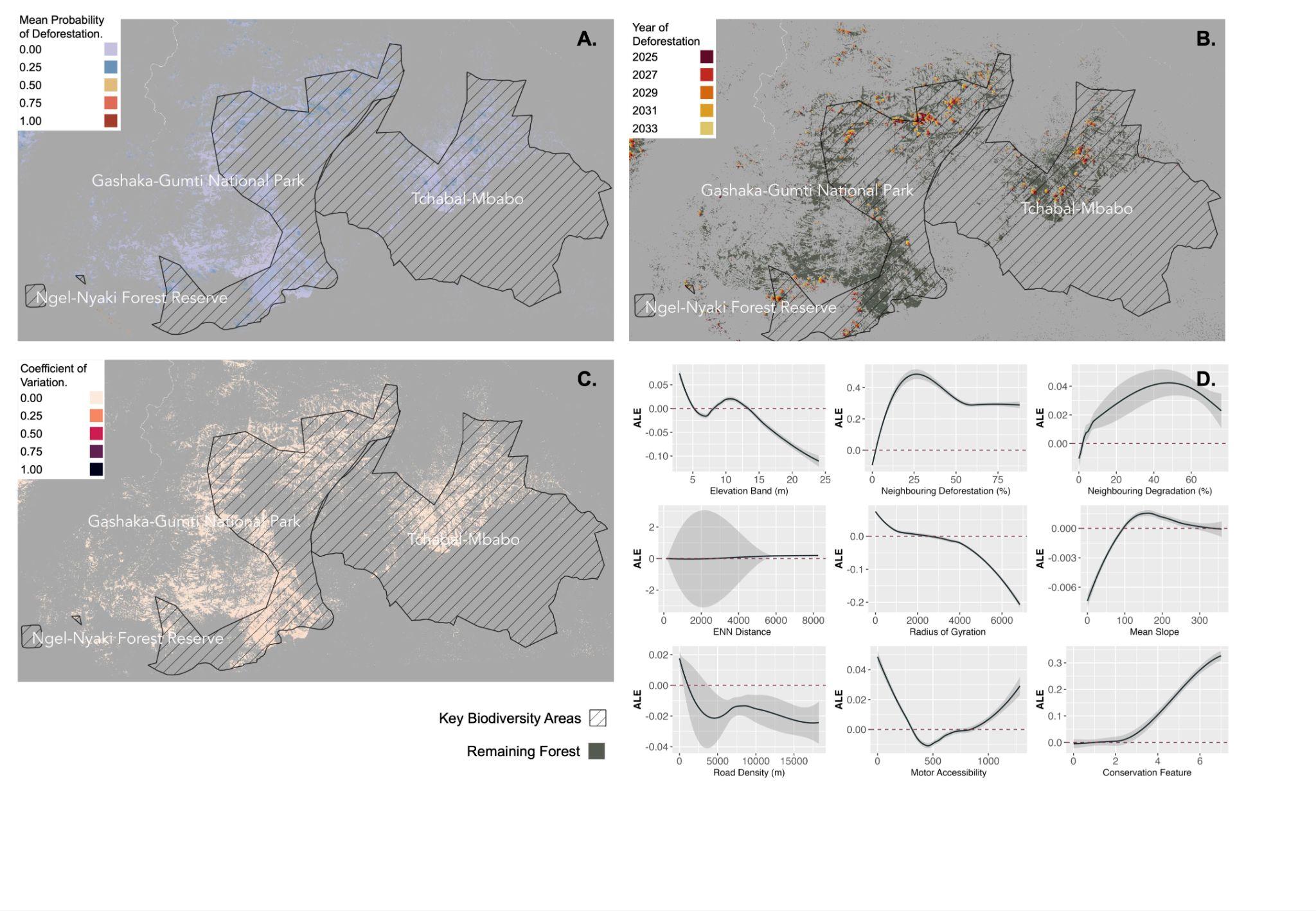

**Figure S3.21 |** Maps across the extent of model group 26, including Key Biodiversity Areas: Gashaka-Gumti NP, Tchabal Mbabo, and Ngel-Nyaki FR for **(A)** Mean probability of deforestation between 2025 and 2033 (inclusive), **(B)** Predicted deforestation event occurrence by year, **(C)** Mean coefficient of variation across model blocks (n = 10), and **(D)** Accumulated local effects plots for explanatory variables.

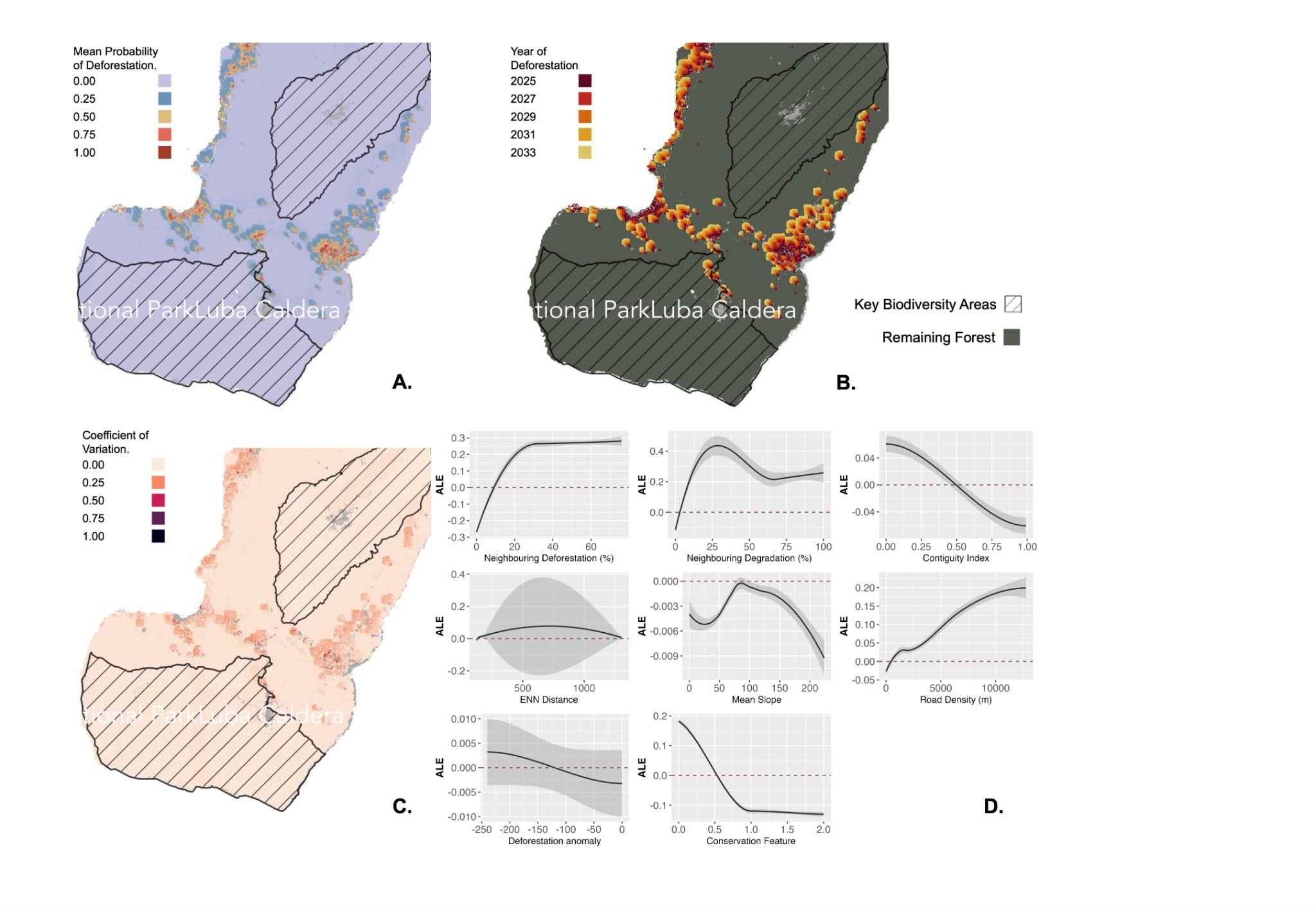

**Figure S3.22 |** Maps across the extent of model group 30, including Key Biodiversity Areas: Luba Caldera Scientific Reserve and Basile Peak NP for **(A)** Mean probability of deforestation between 2025 and 2033 (inclusive), **(B)** Predicted deforestation event occurrence by year, **(C)** Mean coefficient of variation across model blocks (n = 10), and **(D)** Accumulated local effects plots for explanatory variables.
